# Curating MitoCore: A Standardized Small-Scale Human Metabolic Model as Platform for Proteomics Integration and Disease Modeling

**DOI:** 10.64898/2026.06.29.734258

**Authors:** Emanuel Lange, Alejandro Benjamín Reyes Santamaría, Robert Heyer

## Abstract

**Motivation:** Central human metabolism powers cellular processes, yet its dysregulation in disease remains poorly understood. While comprehensive genome-scale metabolic models like Human-GEM are available, their size limits interpretability and computational efficiency. Conversely, the smaller MitoCore model is more manageable but lacks the standardized annotations and curated gene-protein-reaction (GPR) associations necessary for omics integration like protein-constrained modeling. Improving MitoCore’s annotation quality is therefore essential for its use in integrative workflows.

**Results:** We systematically updated MitoCore to enhance compatibility with the protein-constrained modeling framework sMO-MENT. By restructuring legacy annotations and integrating data from Human-GEM and MitoMammal, we increased EC-codes from 354 to 593 and UniProt-annotated genes from 0 to 592. MitoCore captures central metabolic processes, confirmed by mapping its reactions to 51 of 106 metabolic KEGG modules. Integration of thrombocyte proteomics and experimental ATP data for original and curated models showed an increase in mapped proteins (228 to 294) and reactions with kcat values (295 to 310), adding 33 protein-constrained reactions. Consequently, prediction errors for exchange fluxes and ATP production decreased by 19% and 89%, respectively, with 100% of ATP predictions falling within the 95% confidence interval (compared to 16% for the original model). Finally, we implemented a continuous integration/continuous deployment pipeline for automated updates from future Human-GEM releases. These improvements provide a computationally efficient, well-annotated model for studying central metabolism across human cell types.

**Availability and Implementation:** All source code for reproducing results from this paper is available at https://doi.org/10.5281/zenodo.20813825.

## Introduction

Central metabolism takes place in the cytosol and mitochondria of human cells and comprises metabolic reactions that play a central role in converting nutrients to ATP and generating building blocks for biomass. The central metabolism has an important role for tissues and disease conditions and has been subject of investigation to understand metabolism in cardiomyocytes, adipocytes, metabolism in cancer, and immunometabolism (1–4).

In peripheral blood cells such as thrombocytes, metabolism is linked to cellular homeostasis, cellular activation and effector functions including thrombus formation (5, 6). Thrombocytes contain mitochondria and are easily accessible through blood biopsies, which makes them an interesting and accessible subject to investigate mitochondrial dysfunction (5).

Modern analytical methods facilitate the characterization and quantification of healthy and dysregulated metabolism. Involved proteins, metabolites and other molecule classes can be investigated by mass spectrometry (MS)-based omics and nuclear magnetic resonance (NMR). While these methods identify and quantify the involved molecules, other technologies such as ^13^C metabolic flux analysis (7) and Seahorse analysis (8) provide insight into metabolic fluxes. Ultimately, integration of omics and flux data and the corresponding metabolic network is pivotal for data interpretation and mechanistic understanding of dysregulated metabolism.

Genome-scale metabolic reconstructions and its accompanying mathematical implementation, constraint-based models (CBMs), can be utilized for data integration. The default constraint based model compiles a stoichiometric network of metabolic reactions and lower and upper limits (i.e., constraints) for metabolic fluxes (i.e., reaction rates in *mmol/g*_*dw*_*h*) (9). It can be used, for example, to predict flux distributions optimizing a cellular objective (e.g., biomass or energy production) by flux balance analysis (FBA), to explore the metabolic flux variability at the optimum using flux variability analysis (FVA), or the probability distribution of fluxes using flux sampling (9, 10). The space of feasible predictions for flux distributions, i.e., the solution space, can include biologically implausible solutions as cellular resources or regulation of enzymes are not considered by default models. Methods for data integration typically alter model constraints or introduce additional constraints, reducing the solution space, and resulting in model predictions representing the biological contexts more realistically (11). The sMOMENT method, for example, limits the flux of enzymatic reactions by introducing protein resource constraints derived from quantitative proteomics data ^1^ (12).

Prerequisite for data integration is the annotation of genes, reactions and metabolites contained in models with cross-reference to biochemical databases such as UniProt, KEGG, or BiGG (13–15). These annotations allow to uniquely identify model elements, associate measured molecules with model elements or fetch information from databases. Model annotations are supported by the current file format SBML3 (16, 17) and CBM libraries such as COBRApy (18).

High quality constraint-based models applicable to human central metabolism are readily available. Human-GEM is a genome-scale model developed to consolidate multiple genome-scale model lineages into one unified model with standardized identifiers and rich annotations. It has a public version-controlled GitHub repository facilitating community-driven and transparent model development (19). The current version 2.0.0 of Human-GEM contains 12,931 reactions, 8,461 metabolites and 2,848 genes (20) covering many metabolic reactions beyond the central metabolism. However, due to its size, it is difficult to overview and handle manually and requires substantial computational resources for data integration and analysis such as flux sampling.

MitoCore is a less complex metabolic model constituting 555 reactions ^2^, 441 metabolites and 391 genes (1). It represents central pathways for energy, amino and fatty acid metabolism. The main advantage of MitoCore over earlier models and Human-GEM is the explicit modeling of the Proton Motive Force (PMF) by two pseudo metabolites for electrical membrane potential and proton gradient, resulting in more accurate predictions for mitochondrial energy production (1). Recently (2024) the MitoMammal model was published (21), upgrading MitoCore from SBML2 to SBML3, mapping human to murine genes, and introducing the reduction of CoQ by DHODH within the de novo pyrimidine synthesis pathway in form of 5 new reactions. While MitoCore represents central metabolic processes more specifically than Human-GEM, is smaller and therefore feasible for computationally demanding analyses, it lacks standardized annotations preventing automated data integration.

In this paper we aimed to transform MitoCore into a suitable candidate for omics data supported modeling of the human central metabolism. To this end, 1) MitoCore was curated by parsing and cleaning unstructured annotations, adding updated reactions and gene protein reaction (GPR) rules from MitoMammal and enriched the model with annotations from Human-GEM. 2) The coverage of metabolic processes represented in MitoCore and Human-GEM was evaluated by mapping model reactions to KEGG modules aiming to provide a comparative overview of metabolic processes. 3) The influence of our model curation was investigated by integrating a proteomics and Seahorse datasets from resting thrombocytes into different curation states of MitoCore using sMOMENT and performing predictions of metabolic fluxes using flux sampling. 4) Building on the continuous effort of the Human-GEM development, we implemented a continuous integration and continuous deployment (CI/CD) pipeline automatically updating GPRs and annotations of MitoCore with future releases of Human-GEM.

## Methods

The source code for reproducing computational experiments including model curation, proteomics integration, and model analysis are available at https://doi.org/10.5281/zenodo.20813825. All computations were performed on a Windows 10 laptop (Intel i5-1145G7 processor, 16 GB RAM) except for flux sampling (section E), which was performed on a Linux server (AMD EPYC 7543 32-Core processor, 512 GB RAM).

### A. Model Curation

Model curation was conducted in four stages, which included 1) updating and cleaning the original MitoCore model, 2) making MitoCore compatible with proteomics integration, 3) inserting updates introduced in MitoMammal, and 4) inserting annotations and GPR associations from Human-GEM (figure A). After each step, a model was generated, which was used as starting point in the subsequent stage. Additionally, the output models were used to evaluate the impact on information contained in the model, proteomics integration by sMOMENT and model predictions.

In step 1, the original MitoCore model (1) was updated from SBML2 to SBML3 version 1 using COBRApy. Unstructured annotations for genes, reactions, and metabolites - previously stored as strings in the notes object - were parsed and transferred to annotation objects complying with SBML3 specifications. Extracted fields included KEGG compound and BiGG (Recon2) identifiers for metabolites, as well as EC-codes and BiGG (Recon2), Ensembl, HGNC, and KEGG identifiers for reactions and genes. We also cleaned attributes of unusable strings, for example, ensebml identifiers (22) contained entries such as “Unknown”, “N/A”, “Non-Enzymatic”, “Non-enzymatic”. The resulting model is termed “MitoCore Original”.

Step 2 focused on providing the minimum annotations required for the sMOMENT method via the autoPACMEN toolbox (12), specifically EC numbers for reactions, BiGG metabolites, and UniProt accessions for genes. As UniProt accessions were missing, they were retrieved via the HGNC API (23) and added to the model. This version is termed “MitoCore Preliminary”.

**Fig. 1.**
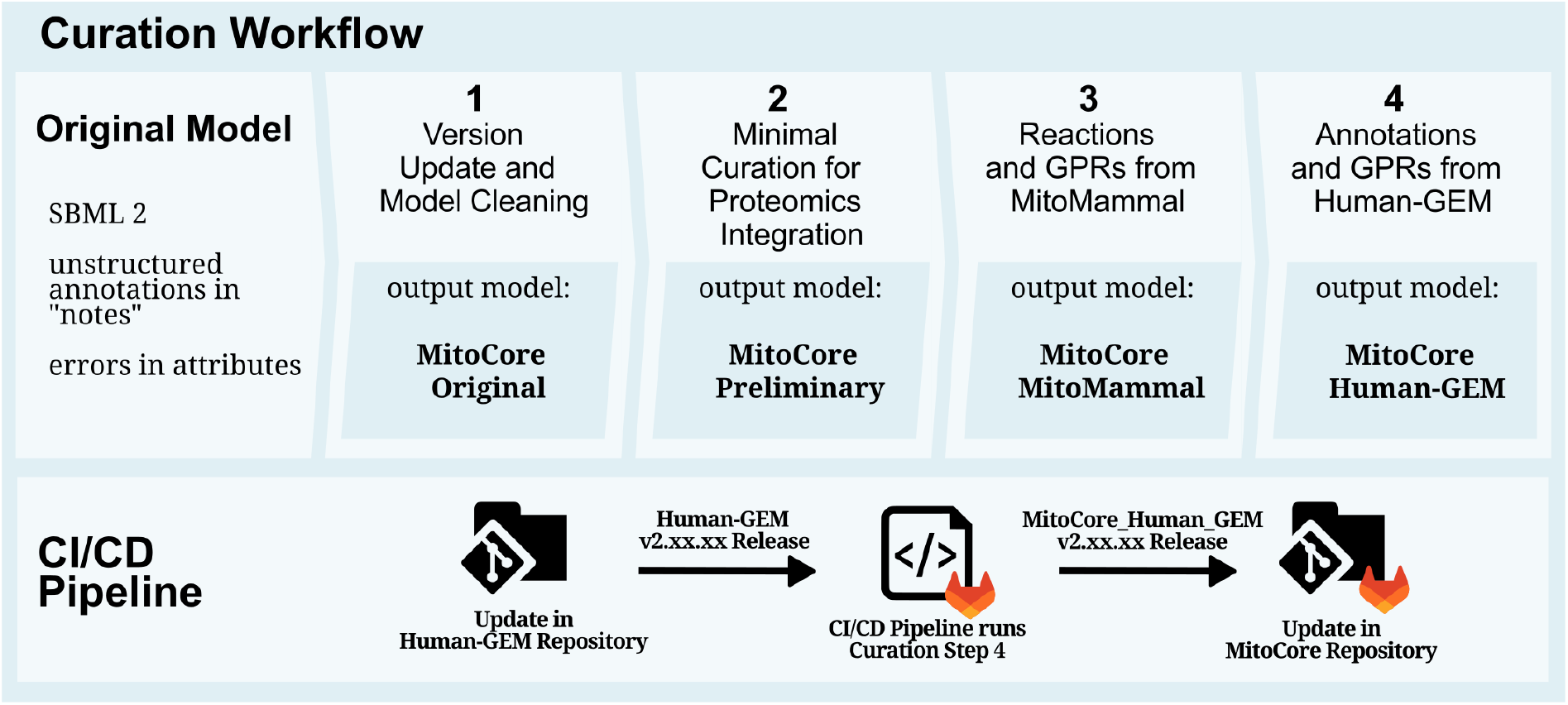
Overview of the curation workflow. Starting with the original MitoCore model in step 1, annotations contained in the notes are parsed into annotation objects. Step 2 introduces the minimally required annotations to make MitoCore ready for the sMOMENT method. Step 3 adds updates introduced to the MitoMammal model. Lastly, reactions, metabolites and genes are mapped to the Human-GEM model, and annotations from Human-GEM are added to MitoCore’s annotations in step 4. For continuous updates of MitoCore annotations, a GitLab CI/CD pipeline was implemented to automatically update MitoCore MitoMammal by running curation step 4 whenever a new version of Human-GEM is released.

In step 3, we incorporated manually curated reactions and modified GPRs from MitoMammal to improve the representation of mitochondrial processes. This included five reactions for CoQ reduction by DHODH, corrected GPRs for electron transport chain (ETC) complexes I and IV, and the addition of the UCP1 gene. This version is termed “MitoCore MitoMammal”. Step 4 aimed to enrich annotations to increase autoPACMEN compatibility and facilitate the mapping of other omics data (e.g., metabolomics via ChEBI identifiers (24)) using Human-GEM as a resource. To enable mapping of reactions between MitoCore and Human-GEM, MetaNetX identifiers (25) were added to MitoCore reactions via BiGG identifiers. Reactions were mapped using a combination of BiGG, KEGG, and MetaNetX to maximize coverage; metabolites were mapped via KEGG, and genes via ensembl identifiers. Annotations from the mapped Human-GEM elements were then transferred to MitoCore. Additionally, MetaNetX identifiers were added to metabolites via KEGG and BiGG identifiers, and MetaNetX (version 4.5) was queried to supplement reaction EC-numbers. Finally, Human-GEM GPR associations were integrated; isoenzymes (genes connected by OR rules) were added if the MitoCore GPR was missing or differed from the Human-GEM isoenzyme. Any new genes identified during this process were added to the model. Manually curated reactions introduced from MitoMammal were assumed to be correct and contained extensive annotations and were therefore protected from any automated updates. The final version is termed “MitoCore Human-GEM”.

### B. KEGG Module Mapping

To analyze the coverage of metabolic pathways by MitoCore and Human-GEM, model reactions were mapped to KEGG modules retrieved by API requests. MitoCore only contains compartments for cytosol, mitochondrion and an external compartment, therefore only reactions from the corresponding compartments in Human-GEM were considered. The coverage was calculated as the fraction of reactions in a KEGG module that could be mapped to model reactions via KEGG reaction identifiers.

### C. Creating sMOMENT Models

As an exemplary use-case, all generated models were parameterized to represent resting thrombocytes assuming maximization of total ATP production (model reaction OF_ATP_MitoCore) as “cellular objective”. We collected publicly available ^13^C data (7), as well as an existing human thrombocyte model (iAT-PLT-636) (26). The constraints of reactions in the “Boundary conditions” group of MitoCore were blocked if no exchange reaction for that metabolite was present in iAT-PLT-636. Furthermore, the maximal uptake or excretion constraints for Glucose, Acetate and Lactate were set to the upper bounds from resting platelets in the ^13^C data. A source for glucose-6-phosphate was introduced to model utilization of glycogen storages. The upper limit for this “glycogen supply” was set to the upper limit determined from the ^13^C analysis. To evaluate the impact of the individual model curation steps, quantitative proteomics data were incorporated into each suitable model version (Preliminary, MitoMammal, Human-GEM) using the autoPACMEN library (12). Essentially, sMOMENT introduces an upper limit to reactions determined from the product of available enzyme concentration and the reaction’s *k*_*cat*_ value (for more information see the original manuscript or supplementary information section 1).

The BiGG (version 1.6) (15), BRENDA (version 2024_1) (27) and SABIO-RK (28) databases were downloaded/queried and parsed as described in the autoPACMEN manual (12). Publicly available proteomics data for platelets (29) were downloaded and transformed from copies/cell to *mmol/g*_*dw*_ utilizing molecular weights queried from UniProt (13) and platelet dry weight (2.13 pg) (30). The cellular protein fraction was calculated by summing individual protein weights based on the proteomics data and dividing by cellular dry weight. The fraction of model included enzymes was calculated by mapping proteins from the data to the model, summing the weight of measured proteins and dividing by the weight of all cellular proteins. The parameters for cellular protein fraction and model included enzymes were passed to autoPACMEN according to its manual. Reactions without *k*_*cat*_s were excluded from sMOMENT to prevent that a default *k*_*cat*_ is introduced, potentially limiting the objective. Candidate reactions for calibration were selected based on a novel algorithm (supplementary information section 1).

For model calibration a publicly available Seahorse dataset was retrieved (8). Scenarios for model calibration were prepared according to autoPACMEN’s manual, setting the target for the objective function to the mean value of total ATP production from the Seahorse data (1.707 *mmol/g*_*dw*_*h*). Calibration was conducted using our version of the “calibrator” (supplementary informtaion section 1).

It is important to note that the integrated datasets originate from distinct experimental conditions. The proteomics data were generated from donor-derived isolated thrombocytes, capturing a physiological state of the proteome (29). Both the ^13^C (7) and Seahorse (8) studies incubated donor-derived thrombocytes in cell-culture media (Tyrode’s and DMEM, respectively). For the final flux measurements, acetate and glucose were supplied as carbon and energy sources during the ^13^C study, whereas the Seahorse assay supplied only glucose and glutamine. Because glutamine was not found to be taken up significantly by thrombocytes (7), its uptake was blocked in our simulations. Consequently, the ATP production derived from the Seahorse data was used to define the baseline, glucose-dependent energy demand (the FBA objective function), while the exchange fluxes from the ^13^C analysis are treated as the broader, physiologically possible boundary constraints.

### D. Analysis of the Solution Space

The solution space was investigated for all models using FVA. autoPACMEN splits reversible reactions with enzyme constraints into forward and reverse reactions creating futile cycles, which can bias FVA results with thermodynamically infeasible solutions. To prevent this bias during FVA, reverse reactions were blocked during maximization of the opposing reaction as done in literature (12). To make all model reactions comparable, every non-protein-constrained reaction was split as well, excluding exchange reactions. For each reaction the flux range was determined as the difference between minimum and maximum flux determined during FVA and summed to obtain a cumulative flux variability (due to splitting, all fluxes ranged between 0 and 1,000 *mmol/g*_*dw*_*h*). Flux ranges for reactions that were split by autoPACMEN were determined from the arm reaction representing the net flux through the original reaction.

### E. Predicting metabolic Fluxes by Flux Sampling

Thrombocytes are anucleate cells without growth, which produce ATP to maintain cellular homeostasis when resting (5). Therefore, we assumed ATP production as cellular objective and that the value measured by Seahorse analysis (1.707 *mmol/g*_*dw*_*h*) represents the average optimal value for resting thrombocytes. Flux sampling was used to explore flux distributions near this measured mean ATP production. We wanted to explore the flux distributions near this assumed optimum and therefore only allowed flux samples that satisfy ATP production that are at least 95% of the measured value by setting a lower limit to the ATP objective function (1.622 *mmol/g*_*dw*_*h*). For flux sampling, metabolic networks were imported to Python using PolyRound (31) and subjected to polytope rounding using the hopsy library (32). The rounded polytopes were sampled using the uniform coordinate hit-and-run proposal (33) implemented in hopsy and configured to create a total of 40,000 samples across 16 Markov chains with a thinning corresponding to the number of reactions times 10. To verify convergence of the sampling algorithm, we verified that the rank-normalized potential scale reduction factors 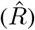 were below 1.01 and the effective sample sizes (ESS) were above 400 per Markov chain as described in literature (34).

### F. CI/CD for Model Updates in GitLab

Git is a version control system for tracking versions of developed software in repositories, which can be made available to collaborators or published through platforms such as GitHub or GitLab (35, 36). GitLab supports CI/CD a concept employed in software development, involving automatic testing of code commits and deploying software automatically via pipelines (37). For automated model updates, a GitLab CI/CD pipeline (38) was implemented, located in a dedicated GitLab repository (https://gitlab.com/mdoa-group/mitocore-human-gem) and scheduled to run every week. The pipeline consists of a build stage, collecting all necessary files, a run stage, executing curation step 4 based on the MitoCore MitoMammal and the latest Human-GEM version, and a deploy stage, running tests, generating reports and creating a tagged commit with the updated model. CI/CD tasks are passed to a GitLab Runner (39), running on a server hosted by the Multidimensional Omics Data Analysis group at the ISAS e.V.

## Results

### G. The Curation Workflow enriches MitoCore with standardized Annotations

The original MitoCore model was published in SBML2 format, which only supports annotations of model elements in the notes object. Technically, these annotations are machine readable and can be used for data integration, but there is no specification how SBML notes should be structured. Therefore the first aim of this work was to cast MitoCore’s annotations into annotation objects supported by SBML3’s specifications and add further ones to make the model compatible with data integration libraries such as autoPACMEN.

Step 1 and 2 were minimal steps required to subject the model to autoPACMEN, which included parsing the required EC numbers and BiGG metabolite identifiers and adding UniProt accessions for every gene (figure 2). Updates from MitoMammal integrated in step 3 increased the total number of reactions, metabolites and genes by 5, 4 and 2, respectively (table 1) and adding 3 EC codes and 12 new isoenzymes to reactions (figure 2). By step 4, additional annotations and GPRs were mined from Human-GEM increasing the number of genes by 208 and the number of genes with UniProt id by 205. EC-codes were increased by 236 and Isoenzymes by 545 (figure 2). Further annotations were added in addition to those mentioned. The overall consistency and annotation content of the models was tested using the MEMOTE test suite (40) yielding an increase of the total score from 35% to 65% from Original to the MitoCore Human-GEM model (figure 2).

**Table 1.** Overview on sizes of the models after the different curation steps and the Human-GEM model.

|  | MitoCore Original/Preliminary | MitoCore MitoMammal | MitoCore Human-GEM | Human-GEM v2.0.0 |
| --- | --- | --- | --- | --- |
| genes | 387 | 389 | 597 | 2,848 |
| reactions | 555 | 560 | 560 | 12,931 |
| metabolites | 441 | 445 | 445 | 8,461 |

**Fig. 2.**
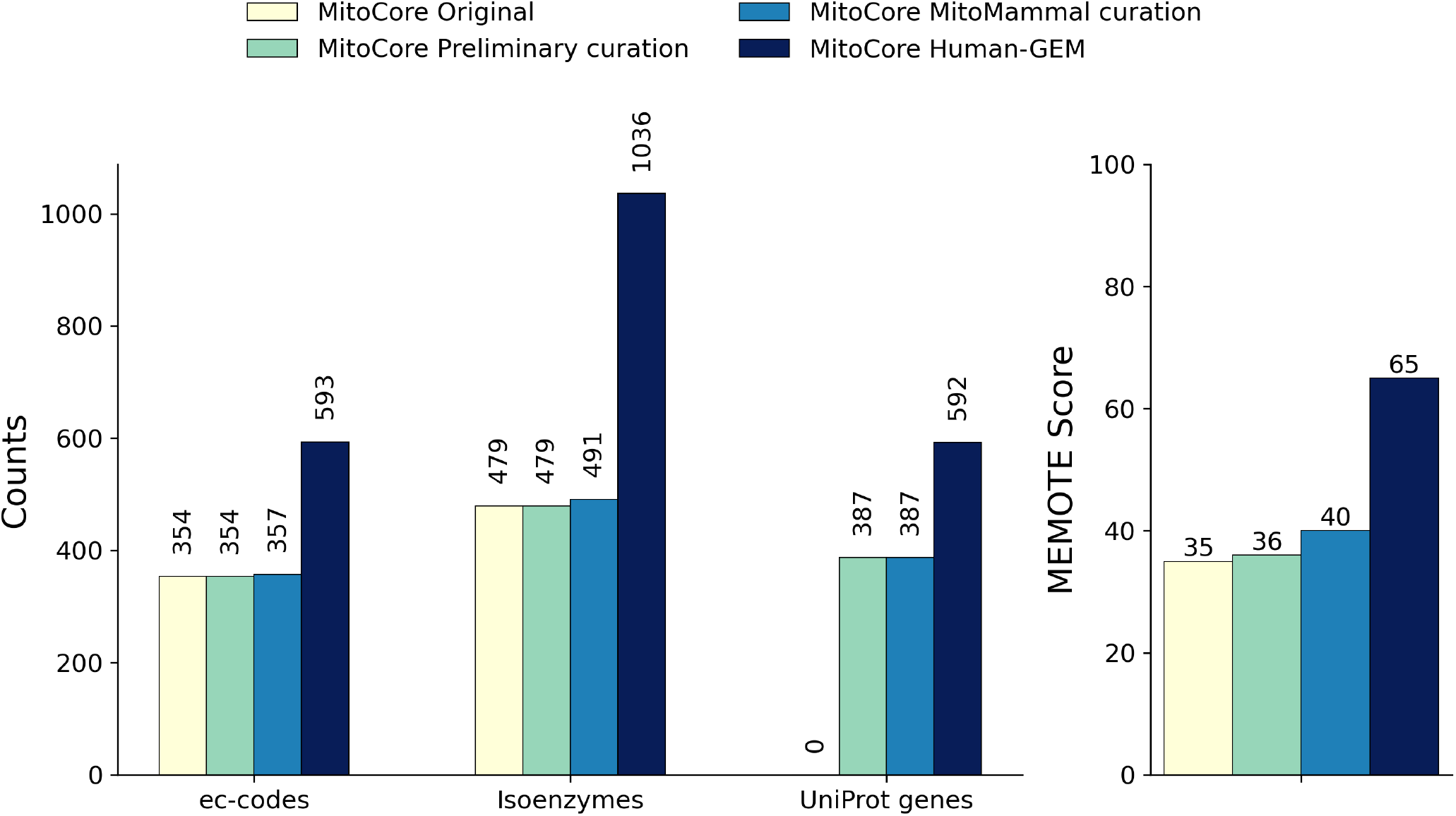
Total counts of EC numbers, isoenzymes and genes with UniProt annotation, as well as the MEMOTE score after each curation step. Shown are annotations with relevance for the autoPACMEN workflow. Isoenzymes represent parts of GPRs that are connected by an OR relationship.

**Fig. 3.**
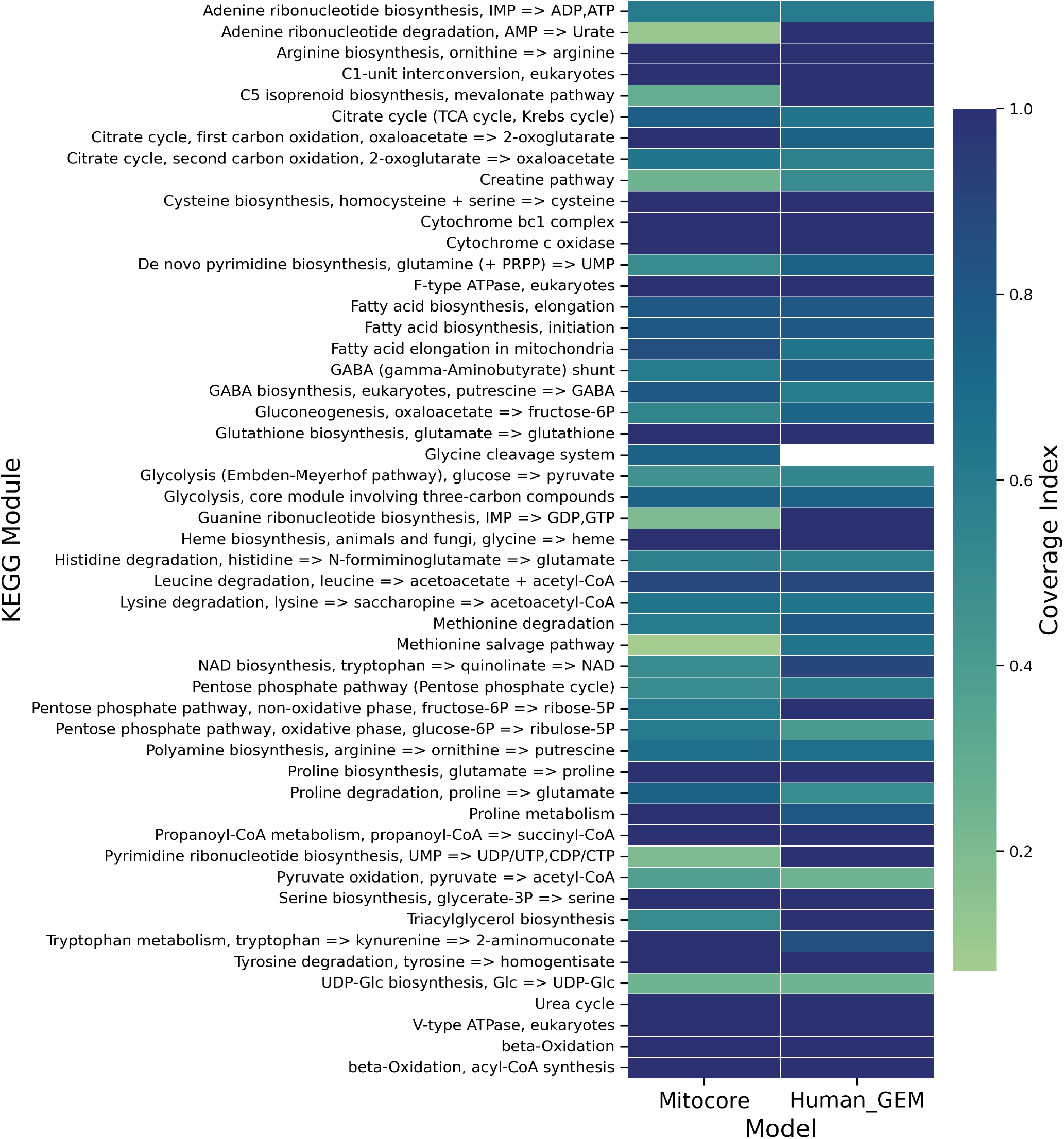
Coverage of KEGG modules by reactions in MitoCore and Human-GEM. The coverage index represents the fraction of reactions in a KEGG module that could be mapped to model reactions. See section 4 in the supplementary information for an overview about all KEGG modules.

For automated updates of MitoCore with information from the latest Human-GEM version, a CI/CD pipeline was implemented in GitLab. The pipeline uses the curated MitoCore MitoMammal model as a starting point and inserts annotations from the latest version of Human-GEM. It includes automated testing to deploy a new MitoCore release whenever annotations are updated. Each deployment generates a detailed report specifying the metabolites, genes, and reactions for which new GPR rules, EC numbers, or UniProt accessions were added, along with a MEMOTE (40) model comparison report to track changes in model quality and coverage.

### H. KEGG Module Mapping supports MitoCore’s Specificity to mitochondrial Processes

MitoCore is a small-scale model focusing on the human central metabolism, while Human-GEM aims to represent all metabolic reactions encoded in the human genome. While it is logical that MitoCore covers fewer metabolic processes than Human-GEM, we wanted to investigate how many elements in MitoCore are contained in Human-GEM and which actual metabolic processes evaluated by the coverage of KEGG modules are included in the model.

Human-GEM covers 100% of all genes, 72.7% of reactions and 94.6% of metabolites contained in MitoCore. 106 human KEGG modules were obtained via API request covering the 555 reactions of MitoCore and the comparable subset of 9,634 Human-GEM reactions (cytosol, mitochondria and extracellular space). The small-scale MitoCore covers 51, while the large-scale Human-GEM covers 105 out of the 106 KEGG modules. There are a few modules uniquely or higher covered by MitoCore, such as modules linked to the TCA cycle, Fatty acid elongation in mitochondria, Glycine cleavage system, and modules linked to Proline and Tryptophan metabolism (see supplementary information section 4).

### I. Standardized Annotations facilitate the sMOMENT Method and impact Model Predictions

Genes, reactions and metabolites contained in metabolic models and their annotations determine which data can be mapped to these model elements and what information can be retrieved from external resources. To investigate this dependency, we parametrized every curation stage of MitoCore to represent resting platelets and integrated proteomics data using the sMOMENT method from the autoPACMEN library.

The original MitoCore model was not applicable to autoPACMEN because the library requires genes to be annotated with UniProt Accessions to request molecular masses from UniProt and map proteins contained in the proteomics data to model genes. After introduction of UniProt Accessions, 228 proteins of 387 with UniProt Accession were mapped to the Preliminary and MitoMammal models and 294 of 597 were mapped to the MitoCore Human-GEM model (Figure 4). AutoPACMEN obtains kcats from the BRENDA and SABIO-RK databases and maps them to reactions annotated with EC-numbers and metabolites with BiGG annotations. For determining the statistics, we counted reactions that had received at least one *k*_*cat*_, but it should be noted that autoPACMEN can assign individual *k*_*cat*_s to forward and reverse reactions. The total amount of reactions with at least one *k*_*cat*_ increased slightly from 295, to 298 to 310 in the Preliminary, MitoMammal and Human-GEM MitoCore models, respectively. Protein constraints are integrated into a reaction if a protein could be mapped to a gene, a molecular mass, and a *k*_*cat*_ is available. Resultingly, The number of reactions receiving a protein constraints amounts to 168, 170, and 201 for Preliminary, MitoMammal and Human-GEM models, respectively.

**Fig. 4.**
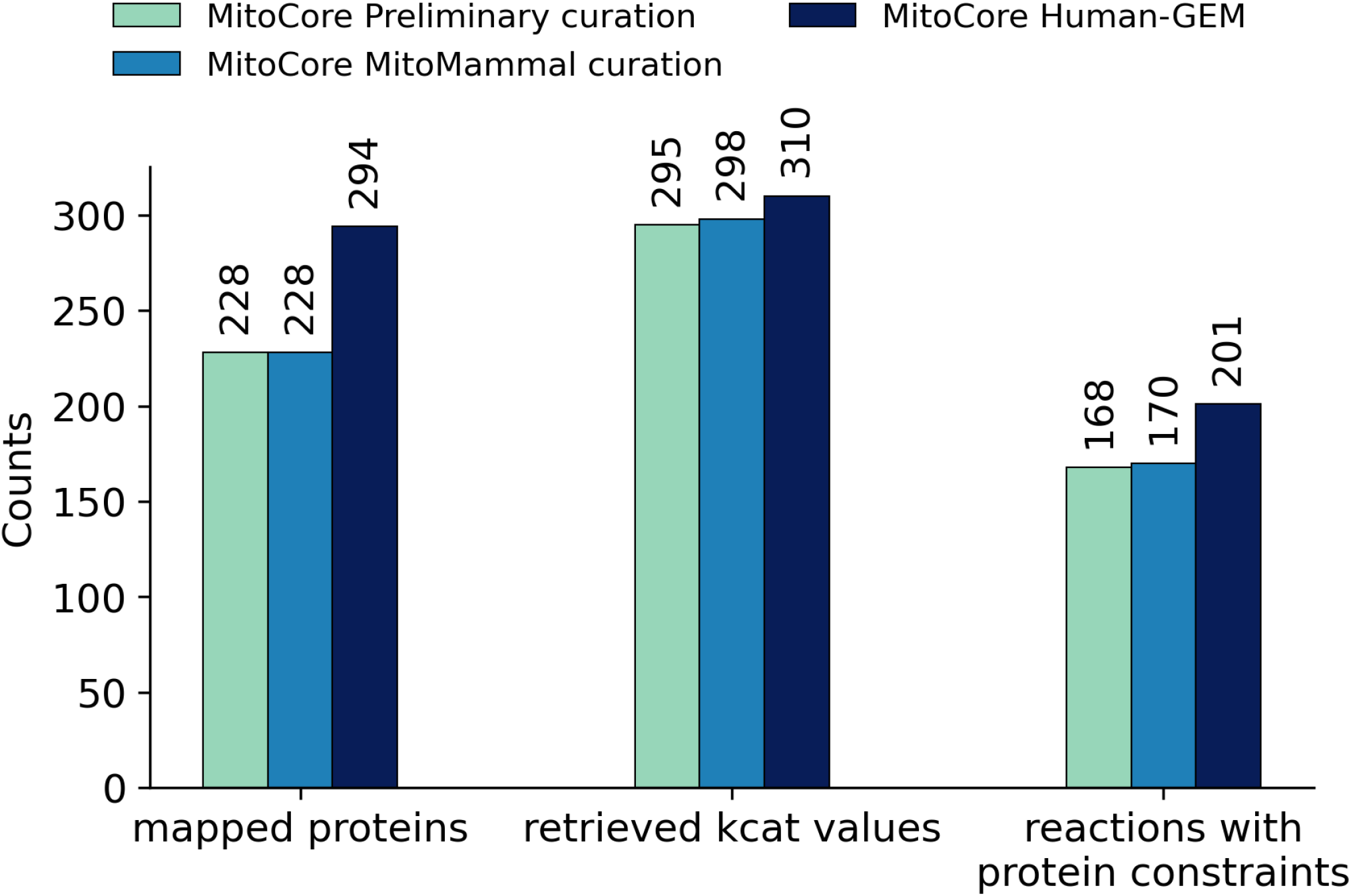
Information retrieved by autoPACMEN for each model applicable to the workflow. Counts represents the absolute number of, e.g., mapped proteins to a model or retrieved *k*^*cat*^s.

Resting platelets require energy for maintaining cellular homeostasis (5), therefore ATP production was used as objective for FBA when running autoPACMEN. The average total ATP production from the Seahorse analysis was set as calibration target. All models with enzyme constraints showed a solution space reduced to approximately 66% of the original model based on the sum of all flux ranges (figure 5 A) and lower average flux ranges of 38 *mmol/g*_*dw*_*h* compared to the original model (58 *mmol/g*_*dw*_*h*). This is also reflected in the IQR limits determined for predicted exchange fluxes and ATP productions (table 2 and 3). To explore metabolic fluxes near the measured mean ATP production (1.707 *mmol/g*_*dw*_*h*), we set a lower bound of 95% of that mean to the ATP objective function and performed flux sampling (figure 5 B; supplementary information, figures S2 to S7). Median predictions from sampling were compared to the means of measurements for exchanged metabolites (glucose, acetate and lactate) and ATP produced by glycolysis and OXPHOS (tables 2 and 3).

**Table 2.** Measured vs. predicted exchange fluxes and prediction errors. Fluxes are represented by their median values in *mmol/g*_*dw*_*h* and corresponding 25% and 75% quantiles of flux sampling distributions in brackets. Quantiles could not be calculated for measuremed fluxes because no standard deviation was provided by (7). The RMSRE (see supplementary information section 2) represents the average relative prediction error of measured fluxes for each model.

| Reaction Name | Measurement | MitoCore Original | MitoCore Preliminary | MitoCore MitoMammal | MitoCore Human-GEM |
| --- | --- | --- | --- | --- | --- |
| glucose uptake | 0.141 | 0.183 (0.185/0.180) | 0.184 (0.185/0.183) | 0.185 (0.185/0.184) | 0.185 (0.186/0.184) |
| acetate uptake | 0.442 | 0.033 (0.046/0.020) | 0.000 (0.001/0.000) | 0.000 (0.001/0.000) | 0.001 (0.002/0.000) |
| lactate secretion | 0.397 | 0.006 (0.003/0.012) | 0.245 (0.241/0.249) | 0.277 (0.273/0.281) | 0.273 (0.269/0.276) |
| RMSRE | - | 0.800 | 0.645 | 0.631 | 0.632 |

**Table 3.** Measured vs. predicted glycolysis and OXPHOS fluxes and prediction errors. Fluxes are represented by their median values in *mmol/g*_*dw*_*h* and corresponding 25% and 75% quantiles of measurements and flux sampling distributions in brackets. The RMSRE represents the average relative prediction error of measured fluxes for each model. The MCP represents the average fraction of flux samples inside the 95% confidence intervals of the measurements (see supplementary information section 2).

| Reaction Name | Measurement | MitoCore Original | MitoCore Preliminary | MitoCore MitoMammal | MitoCore Human-GEM |
| --- | --- | --- | --- | --- | --- |
| glycolysis ATP | 0.411<br>(0.299/0.523) | 0.353<br>(0.340/0.364) | 0.371<br>(0.369/0.374) | 0.372<br>(0.370/0.374) | 0.372<br>(0.371/0.373) |
| OXPHOS ATP | 1.296<br>(1.024/1.568) | 2.785<br>(2.524/3.071) | 1.175<br>(1.170/1.179) | 1.200<br>(1.197/1.202) | 1.183<br>(1.183/1.184) |
| RMSRE | - | 0.819 | 0.095 | 0.085 | 0.091 |
| MCP | - | 0.163 | 0.771 | 0.996 | 1.000 |
| glycolytic index | 0.241<br>(0.177/0.303) | 0.112<br>(0.102/0.123) | 0.240<br>(0.239/0.242) | 0.237<br>(0.236/0.238) | 0.239<br>(0.238/0.240) |

**Fig. 5.**
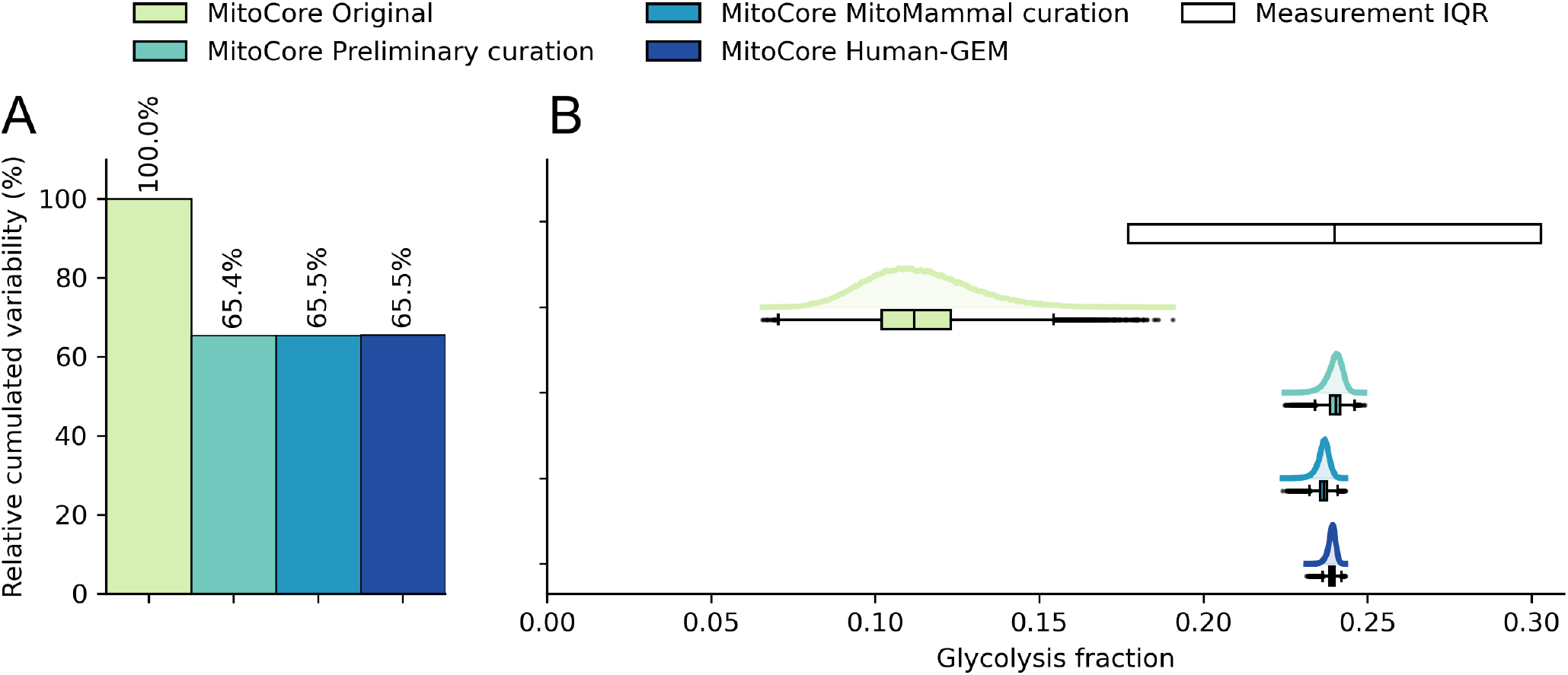
Model analysis. A - Relative size of the solution space of each model compared to the original model. The solution space is represented by the sum of flux ranges of each reaction. B - Measured glycolysis fraction (the box borders represent the estimated IQR and the vertical line in its center the median glycolysis fraction) and predicted flux distributions by flux sampling. IQR of the measured glycolysis fraction was estimated via error propagation of the standard deviation of net fluxes (see supplementary information). Shown flux sampling distributions support at least 95% of the FBA optimum of total ATP production.

All models predicted the uptake of glucose, while only protein-constrained models predicted the secretion of lactate. Acetate was measured to be taken up, but only the MitoCore Original model predicted a considerable uptake flux, which was one magnitude lower than the measurement (table 2). The total relative prediction errors reveal that the protein-constrained MitoCore MitoMammal and Human-GEM models achieved the most accurate predictions (RMSRE 0.631 and 0.632, respectively), followed by the Preliminary (RMSRE 0.645) and non-protein-constrained Original model (RMSRE 0.800).

Analysis of platelet energetics has been considered as a proxy to investigate mitochondrial dysfunction (5, 8) and therefore we also tested predictions of ATP produced from the two main energy delivering processes, glycolysis and OXPHOS. For the absolute ATP production fluxes, all protein-constrained models had similar relative prediction errors (RMSRE 0.085 to 0.095), which were one magnitude lower compared to the Original model (RMSRE 0.819). As the Seahorse data (8) contained information about the cohort size and a standard deviation, we could estimate 95% confidence intervals for the data. To account for the spread of flux sampling distributions, we evaluated the fraction of flux samples falling into this biologically feasible range (MCP, see supplementary information section 2). More than 99% of flux samples from MitoCore MitoMammal and Human-GEM are located in this range (MCP 0.996 and 1.000, respectively), while MitoCore Preliminary and the Original model reached 77% (MCP 0.771) and 16% (MCp 0.163), respectively.

Additionally, we investigated the glycolytic index, which represents the fraction of total ATP produced by glycolysis and is used as an indicator for metabolic shifts (41). Seahorse data represents a glycolytic index of 0.241, which could be represented most accurately by the protein-constrained models (0.237 to 0.240) (figure 5 B).

## Discussion

This study provides an extensive update to MitoCore and its annotations, demonstrating the applicability of state-of-the-art constraint-based modeling methods such as sMOMENT and flux sampling. We focused on MitoCore to create a generalized model for investigating central metabolism across different cell types. Despite its outdated format, MitoCore offers an explicit representation of mitochondrial processes (e.g., PMF), existing annotations, and recent updates from the MitoMammal study (21). Such a generalized model can serve as a template for condition-specific models through data integration, which requires standardized annotations for compatibility with pipelines like autoPACMEN.

We utilized Human-GEM as a template due to its accessibility via GitHub and regular updates. This allows for the use of CI/CD pipelines, a method in software developmentpreviously used with the GECKO toolbox to generate protein-constrained models from Human-GEM (42). Similarly, we implemented a CI/CD pipeline triggered by new Human-GEM releases to automatically update MitoCore’s annotations, facilitating ongoing maintenance.

Our sequential curation workflow enabled precise tracking of changes, prioritizing the generation of models with reliable annotations and GPRs before integrating Human-GEM data. While Human-GEM is extensive, its size makes it prone to errors (21), as seen in its GitHub issue tracker (https://github.com/SysBioChalmers/Human-GEM). For example, a minor error in the GPR of the ADP forming reaction for succinyl coenzyme A synthetase (SUCOASm) propagated from Human-GEM to MitoCore. MitoCore’s GPR for this reaction only contains a gene for the ADP forming subunit (ENSG00000136143), while Human-GEM contains a gene of the GDP forming subunit (ENSG00000163541, reaction MAR04152) resulting in a merged GPR including both genes in MitoCore Human-GEM. onsequently, MitoCore Human-GEM may be unsuitable for applications requiring high mapping specificity, though such errors are decreasing with regular updates of Human-GEM (20).

While originally parameterized for myocardial metabolism, MitoCore is a generalized model for human central metabolism and has been applied to other tissues, such as brown adipose tissue (21). As it only represents the central metabolism, it is important to know whether metabolic processes are covered by MitoCore before parameterizing it to a new cell type. To this end we provided a mapping of metabolic reactions in MitoCore and Human-GEM to KEGG modules. As expected, Human-GEM covers nearly all human metabolism KEGG modules except the Glycine cleavage system. Although Human-GEM contains reactions for this system (e.g., MAR08433, MAR08434, MAR06409), they lacked correct KEGG identifiers. While MitoCore covers essential central metabolism modules, some specific ones, such as Catecholamine biosynthesis (M00042), are excluded. Thus, our mapping serves as a lookup to verify the presence of required modules in MitoCore.

To evaluate the impact of our curation, we applied the sMOMENT method to human thrombocytes. These anucleated cells, derived from megakaryocyte fragmentation, are primarily responsible for hemostasis but also perform immune regulatory functions and contribute to disease pathogenesis (5, 10). Thrombocytes are equipped with RNA for protein synthesis and contain organelles, including 5-8 metabolically active mitochondria per cell (5, 10). Because mitochondrial dysfunction is linked to various diseases, thrombocytes are proposed as an accessible target for investigating bioenergetics (5). Thrombocytes were also chosen due to the availability of quantitative proteomics data for protein constraints (29), Seahorse data for calibrating ATP production, ^13^C data (7), and an existing platelet model (26) for parameterization and validation.

We applied autoPACMEN to the models containing the required annotations. While curation step 2 provided the essential UniProt Accessions, extending model genes via Human-GEM increased the number of mappable proteins by 66 (figure 4). The increase in retrieved *k*_*cat*_ values was smaller, despite total EC-codes increasing from 357 to 593 in step 4. This can be explained by the number of reactions with at least one EC-code, which only increased by 10 (from 321 to 331). The total amount of reactions with integrated protein constraints results from the mapped proteins and retrieved *k*_*cat*_ values. The total number of reactions with integrated protein constraints is lower than the number of mapped proteins because multiple proteins can be assigned to one reaction, and autoPACMEN requires all enzyme subunits to be measured. These metrics demonstrate that our curation improved mapped data, retrieved parameters, and integrated constraints.

To determine if the additional constraints impacted model predictions, we implemented a new algorithm to select candidate reactions for calibration and modified the calibrator to be compatible with individually measured protein pools (supplementary information section 1).

FVA analysis showed that protein constraints reduced the solution space as expected. However, the solution spaces for the protein-constrained models were nearly equal in size, indicating that additional constraints in the MitoCore Human-GEM model had a negligible effect. Similarly, protein constraints influenced flux sampling predictions more significantly than additional annotation from the curation steps, though curation (specifically step 3’s MitoMammal updates) had a slight effect. The MitoCore MitoMammal protein-constrained model yielded the lowest overall prediction errors. Contrary to our expectations, integrating Human-GEM information did not further reduce errors. This limited impact may be due to a conserved rate-limiting reaction shared across all protein-constrained models. Sensitivity analysis (supplementary information section 1) revealed that the FBA objective in all models was most sensitive to the *k*_*cat*_ of the ATP-ase reaction (CV_MitoCore, data not shown), suggesting that this reaction’s constraint limits total ATP production.

The ^13^C study indicated that thrombocytes rely primarily on glucose and acetate as carbon sources and secrete lactate (7). The authors also observed metabolic fragmentation in resting platelets where glucose is metabolized via pyruvate into lactate, while acetate fuels the TCA cycle via acetyl-CoA. The conversion of pyruvate from glycolysis to acetyl-CoA, catalyzed by mitochondrial pyruvate dehydrogenase (PDHm reaction) (43), was found to be inactive (7). All protein-constrained models correctly predicted glucose uptake and lactate secretion (Table 2), but none predicted acetate uptake. As mentioned in the methods section (section C), the datasets originated from different experimental conditions, which could be an explanation for this discrepancy. We assume that with acetate and glucose supply (as in the ^13^C analysis), thrombocytes could generate a higher total ATP production rate compared to using only glucose as in the Seahorse analysis. By calibrating our protein-constrained models to this presumably lower ATP production, the model’s protein resource limits for glucose uptake and processing are not reached and acetate uptake is not included in the optimal flux distribution from FBA. This effect could be compounded by the model’s handling of the PDHm reaction (Supplementary Figure S7), fueling the TCA cycle via pyruvate-derived acetyl-CoA. PDH enzyme activity is regulated by phosphorylation, highlighting a case of post-translational modification altering activity without changing protein abundance (43). Such enzyme regulations are not captured by our modeling approach, which considers only total protein abundances.

ATP production predicted for glycolysis and OXPHOS on the other hand were representative of the measured values from Seahorse. The glycolytic index, which represents the glycolytic fraction of total ATP production, indicates whether cells prefer glycolytic or oxidative metabolism (41). A chronic increase in the glycolytic index is, for example, indicative for the Warburg effect occuring in cancer cells (41). Reported indices for thrombocytes vary from balanced use of glycolysis and OXPHOS (5, 7) to a more oxidative metabolism (8). Our models, calibrated using data from young adult thrombocytes (8), consistently predicted a preference for oxidative metabolism.

The integration of data in this study was inherently constrained by the public availability of exchange and ATP production fluxes. Specifically, the absence of donor-specific metadata in the ^13^C dataset precluded the derivation of 95% confidence intervals for error evaluation (Supplementary Information Section 2). Furthermore, as previously discussed, integrating datasets from disparate experimental conditions introduces inherent biological discrepancies. While the current approach effectively benchmarks the impact of model curation, future studies will benefit from datasets derived from a matched donor cohort under identical conditions. In addition, experimental limitations must be considered. For instance, Seahorse analysis relies on simplifying assumptions regarding media molecule ratios and is sensitive to cell treatment (e.g., immediate measurement vs. in vitro culture) (44). While measuring absolute physiological ATP production accurately is challenging, capturing qualitative shifts in ATP production or the glycolytic index between conditions can still yield qualitatively accurate model predictions.

Furthermore, we recommend calibrating models across multiple conditions. While our protein-constrained models represent their specific calibration condition well, their ability to generalize to activated or dysregulated metabolism is unclear. AutoPACMEN supports multi-scenario calibration, which could provide more reliable *k*_*cat*_ parameters. Since thrombocytes can synthesize proteins from mRNA (45), condition-dependent proteome changes could be integrated by using a “consensus” proteomics dataset (maximum measured protein resources across conditions) or by swapping protein constraints while keeping *k*_*cat*_ values fixed. The latter requires that enzyme activity is mostly dependent on enzyme concentration rather than pos-translational regulation, meaning that *k*_*cat*_s are approximately constant over conditions. However, it cannot be excluded that post-translational regulations occur, as seen with PDHm (43, 46).

We employed the sMOMENT method for its accurate quantitative flux predictions (12). While the GECKO toolbox (47) offers a similar approach, we chose autoPACMEN for its *k*_*cat*_ calibrator and because its implemented in Python for the largest part. A drawback of both is their dependence on proprietary MATLAB licenses. Python-based libraries such as geckopy (48) could serve as an alternative. Another advantage of protein-constrained models is their compatibility to widespread methods such as FBA. An alternative method for proteomics integration is E-Flux, which has been applied in the study of MitoMammal (21). E-flux scales the upper bounds for reactions based on transcriptomics or proteomics data and is compatible to absolute and relatively quantified proteins. While it requires less data than autoPACMEN, it can only provide qualitative predictions. Finally, the curated MitoCore Human-GEM model includes metabolite cross-references (e.g., ChEBI (24), PubChem (49)), making it compatible with metabolomics data mapping.

## Supplementary Note 1: Competing interests

No competing interest is declared.

## Supplementary Note 2: Author contributions statement

- Lange, E. — Conceptualization (lead), Methodology (lead), Software (lead), Validation (lead), Formal analysis (lead), Investigation (lead), Data curation (lead), Writing — original draft (lead), Writing — review & editing (lead), Visualization (lead), Supervision (equal).
- Reyes Santamaria, A.B. — Conceptualization (supporting), Methodology (supporting), Software (supporting), Validation (supporting), Formal analysis (supporting), Investigation (supporting), Data curation (supporting), Writing — original draft (supporting), Writing — review & editing (supporting), Visualization (supporting).
- Heyer, R. — Conceptualization (supporting), Resources (lead), Supervision (equal), Project administration (lead), Funding acquisition (lead).

## ACKNOWLEDGEMENTS

All authors acknowledge the support by the “Ministerium für Kultur und Wissenschaft des Landes Nordrhein-Westfalen” and “Der Regierende Bürgermeister von Berlin, Senatskanzlei Wissenschaft und Forschung”. We thank all co-workers and friends for discussions and feedback, which helped to bring this project to its final form.

## Footnotes

1 In this paper we refer to models containing protein-resource constraints as protein-constrained models.

2 This number represents the model’s contents after transfer to SBML3. In its original version in SBML2, exchange reactions are not explicitly included.

## Notes

https://doi.org/10.5281/zenodo.20813825

https://gitlab.com/mdoa-group/mitocore-human-gem

### Competing Interest Statement

The authors have declared no competing interest.

### Summary of Updates

Corrections of results in abstract: 33 new protein constraints were added and the prediction error decreased by 89% compared to the original model.

## Bibliography

1. Anthony C. Smith, Filmon Eyassu, Jean-Pierre Mazat, and Alan J. Robinson. Mitocore: a curated constraint-based model for simulating human central metabolism. BMC Systems Biology, 11(1), November 2017. ISSN 1752-0509. doi: 10.1186/s12918-017-0500-7.

2. Marzia Di Filippo, Dario Pescini, Bruno Giovanni Galuzzi, Marcella Bonanomi, Daniela Gaglio, Eleonora Mangano, Clarissa Consolandi, Lilia Alberghina, Marco Vanoni, and Chiara Damiani. Integrate: Model-based multi-omics data integration to characterize multi-level metabolic regulation. PLOS Computational Biology, 18(2):e1009337, February 2022. ISSN 1553-7358. doi: 10.1371/journal.pcbi.1009337.

3. Ziming Cao, Meng Zhao, Hao Sun, Liang Hu, Yunfeng Chen, and Zhichao Fan. Roles of mitochondria in neutrophils. Frontiers in Immunology, 13, August 2022. ISSN 1664-3224. doi: 10.3389/fimmu.2022.934444.

4. Hui Liu, Shuo Wang, Jianhua Wang, Xin Guo, Yujing Song, Kun Fu, Zhenjie Gao, Danfeng Liu, Wei He, and Lei-Lei Yang. Energy metabolism in health and diseases. Signal Transduction and Targeted Therapy, 10(1), February 2025. ISSN 2059-3635. doi: 10.1038/s41392-025-02141-x.

5. Mia S. Wilkinson and Kimberly J. Dunham-Snary. Blood-based bioenergetics: a liquid biopsy of mitochondrial dysfunction in disease. Trends in Endocrinology amp; Metabolism, 34(9):554–570, September 2023. ISSN 1043-2760. doi: 10.1016/j.tem.2023.06.004.

6. Paresh P. Kulkarni, Mohammad Ekhlak, and Debabrata Dash. Energy metabolism in platelets fuels thrombus formation: Halting the thrombosis engine with small-molecule modulators of platelet metabolism. Metabolism, 145:155596, August 2023. ISSN 0026-0495. doi: 10.1016/j.metabol.2023.155596.

7. Cara L. Sake, Alexander J. Metcalf, Michelle Meagher, Jorge Di Paola, Keith B. Neeves, and Nanette R. Boyle. Isotopically nonstationary 13c metabolic flux analysis in resting and activated human platelets. Metabolic Engineering, 69:313–322, January 2022. ISSN 1096-7176. doi: 10.1016/j.ymben.2021.12.007.

8. J JEDLIČ KA, R Kunc, and J KuncovÁ. Mitochondrial respiration of human platelets in young adult and advanced age – seahorse or o2k? Physiological Research, pages S369–S379, December 2021. ISSN 0862-8408. doi: 10.33549/physiolres.934812.

9. Aarash Bordbar, Jonathan M. Monk, Zachary A. King, and Bernhard O. Palsson. Constraint-based models predict metabolic and associated cellular functions. Nature Reviews Genetics, 15(2): 107–120, January 2014. ISSN 1471-0064. doi: 10.1038/nrg3643.

10. Helena A. Herrmann, Beth C. Dyson, Lucy Vass, Giles N. Johnson, and Jean-Marc Schwartz. Flux sampling is a powerful tool to study metabolism under changing environmental conditions. npj Systems Biology and Applications, 5(1), September 2019. ISSN 2056-7189. doi: 10.1038/s41540-019-0109-0.

11. Nathan E. Lewis, Harish Nagarajan, and Bernhard O. Palsson. Constraining the metabolic genotype–phenotype relationship using a phylogeny of in silico methods. Nature Reviews Microbiology, 10(4):291–305, February 2012. ISSN 1740-1534. doi: 10.1038/nrmicro2737.

12. Pavlos Stephanos Bekiaris and Steffen Klamt. Automatic construction of metabolic models with enzyme constraints. BMC Bioinformatics, 21(1), January 2020. ISSN 1471-2105. doi: 10.1186/s12859-019-3329-9.

13. Alex Bateman, Maria-Jesus Martin, Sandra Orchard, Michele Magrane, Aduragbemi Adesina, Shadab Ahmad, Emily H Bowler-Barnett, Hema Bye-A-Jee, David Carpentier, Paul Denny, Jun Fan, Penelope Garmiri, Leonardo Jose da Costa Gonzales, Abdulrahman Hussein, Alexandr Ignatchenko, Giuseppe Insana, Rizwan Ishtiaq, Vishal Joshi, Dushyanth Jyothi, Swaathi Kandasaamy, Antonia Lock, Aurelien Luciani, Jie Luo, Yvonne Lussi, Juan Sebastian Martinez Marin, Pedro Raposo, Daniel L Rice, Rafael Santos, Elena Speretta, James Stephenson, Prabhat Totoo, Nidhi Tyagi, Nadya Urakova, Preethi Vasudev, Kate Warner, Supun Wijerathne, Conny Wing-Heng Yu, Rossana Zaru, Alan J Bridge, Lucila Aimo, Ghislaine Argoud-Puy, Andrea H Auchincloss, Kristian B Axelsen, Parit Bansal, Delphine Baratin, Teresa M Batista Neto, Marie-Claude Blatter, Jerven T Bolleman, Emmanuel Boutet, Lionel Breuza, Blanca Cabrera Gil, Cristina Casals-Casas, Kamal Chikh Echioukh, Elisabeth Coudert, Beatrice Cuche, Edouard de Castro, Anne Estreicher, Maria L Famiglietti, Marc Feuermann, Elisabeth Gasteiger, Pascale Gaudet, Sebastien Gehant, Vivienne Gerritsen, Arnaud Gos, Nadine Gruaz, Chantal Hulo, Nevila Hyka-Nouspikel, Florence Jungo, Arnaud Kerhornou, Philippe Le Mercier, Damien Lieberherr, Patrick Masson, Anne Morgat, Salvo Paesano, Ivo Pedruzzi, Sandrine Pilbout, Lucille Pourcel, Sylvain Poux, Monica Pozzato, Manuela Pruess, Nicole Redaschi, Catherine Rivoire, Christian J A Sigrist, Karin Sonesson, Shyamala Sundaram, Anastasia Sveshnikova, Cathy H Wu, Cecilia N Arighi, Chuming Chen, Yongxing Chen, Hongzhan Huang, Kati Laiho, Minna Lehvaslaiho, Peter McGarvey, Darren A Natale, Karen Ross,R R Vinayaka, Yuqi Wang, and Jian Zhang. Uniprot: the universal protein knowledgebase in 2025. Nucleic Acids Research, 53(D1):D609–D617, November 2024. ISSN 1362-4962. doi: 10.1093/nar/gkae1010.

14. Minoru Kanehisa, Miho Furumichi, Yoko Sato, Yuriko Matsuura, and Mari Ishiguro-Watanabe. Kegg: biological systems database as a model of the real world. Nucleic Acids Research, 53(D1): D672–D677, October 2024. ISSN 1362-4962. doi: 10.1093/nar/gkae909.

15. Zachary A. King, Justin Lu, Andreas Dräger, Philip Miller, Stephen Federowicz, Joshua A. Lerman, Ali Ebrahim, Bernhard O. Palsson, and Nathan E. Lewis. Bigg models: A platform for integrating, standardizing and sharing genome-scale models. Nucleic Acids Research, 44(D1):D515–D522, October 2015. ISSN 1362-4962. doi: 10.1093/nar/gkv1049.

16. N. Juty, N. Le Novere, and C. Laibe. Identifiers.org and miriam registry: community resources to provide persistent identification. Nucleic Acids Research, 40(D1):D580–D586, December 2011. ISSN 1362-4962. doi: 10.1093/nar/gkr1097.

17. Sarah M Keating, Dagmar Waltemath, Matthias König, Fengkai Zhang, Andreas Dräger, Claudine Chaouiya, Frank T Bergmann, Andrew Finney, Colin S Gillespie, Tomáš Helikar, Stefan Hoops, Rahuman S Malik-Sheriff, Stuart L Moodie, Ion I Moraru, Chris J Myers, Aurélien Naldi, Brett G Olivier, Sven Sahle, James C Schaff, Lucian P Smith, Maciej J Swat, Denis Thieffry, Leandro Watanabe, Darren J Wilkinson, Michael L Blinov, Kimberly Begley, James R Faeder, Harold F Gómez, Thomas M Hamm, Yuichiro Inagaki, Wolfram Liebermeister, Allyson L Lister, Daniel Lucio, Eric Mjolsness, Carole J Proctor, Karthik Raman, Nicolas Rodriguez, Clifford A Shaffer, Bruce E Shapiro, Joerg Stelling, Neil Swainston, Naoki Tanimura, John Wagner, Martin Meier-Schellersheim, Herbert M Sauro, Bernhard Palsson, Hamid Bolouri, Hiroaki Kitano, Akira Funahashi, Henning Hermjakob, John C Doyle, Michael Hucka, Richard R Adams, Nicholas A Allen, Bastian R Angermann, Marco Antoniotti, Gary D Bader, Jan Červený, Mélanie Courtot, Chris D Cox, Piero Dalle Pezze, Emek Demir, William S Denney, Harish Dharuri, Julien Dorier, Dirk Drasdo, Ali Ebrahim, Johannes Eichner, Johan Elf, Lukas Endler, Chris T Evelo, Christoph Flamm, Ronan MT Fleming, Martina Fröhlich, Mihai Glont, Emanuel Gonçalves, Martin Golebiewski, Hovakim Grabski, Alex Gutteridge, Damon Hachmeister, Leonard A Harris, Benjamin D Heavner, Ron Henkel, William S Hlavacek, Bin Hu, Daniel R Hyduke, Hidde de Jong, Nick Juty, Peter D Karp, Jonathan R Karr, Douglas B Kell, Roland Keller, Ilya Kiselev, Steffen Klamt, Edda Klipp, Christian Knüpfer, Fedor Kolpakov, Falko Krause, Martina Kutmon, Camille Laibe, Conor Lawless, Lu Li, Leslie M Loew, Rainer Machne, Yukiko Matsuoka, Pedro Mendes, Huaiyu Mi, Florian Mittag, Pedro T Monteiro, Kedar Nath Natarajan, Poul MF Nielsen, Tramy Nguyen, Alida Palmisano, Jean-Baptiste Pettit, Thomas Pfau, Robert D Phair, Tomas Radivoyevitch, Johann M Rohwer, Oliver A Ruebenacker, Julio Saez-Rodriguez, Martin Scharm, Henning Schmidt, Falk Schreiber, Michael Schubert, Roman Schulte, Stuart C Sealfon, Kieran Smallbone, Sylvain Soliman, Melanie I Stefan, Devin P Sullivan, Koichi Takahashi, Bas Teusink, David Tolnay, Ibrahim Vazirabad, Axel von Kamp, Ulrike Wittig, Clemens Wrzodek, Finja Wrzodek, Ioannis Xenarios, Anna Zhukova, and Jeremy Zucker. <scp>sbml</scp> level 3: an extensible format for the exchange and reuse of biological models. Molecular Systems Biology, 16(8), August 2020. ISSN 1744-4292. doi: 10.15252/msb.20199110.

18. Ali Ebrahim, Joshua A Lerman, Bernhard O Palsson, and Daniel R Hyduke. Cobrapy: Constraints-based reconstruction and analysis for python. BMC Systems Biology, 7(1), August 2013. ISSN 1752-0509. doi: 10.1186/1752-0509-7-74.

19. Jonathan L. Robinson, Pınar Kocabaş, Hao Wang, Pierre-Etienne Cholley, Daniel Cook, Avlant Nilsson, Mihail Anton, Raphael Ferreira, Iván Domenzain, Virinchi Billa, Angelo Limeta, Alex Hedin, Johan Gustafsson, Eduard J. Kerkhoven, L. Thomas Svensson, Bernhard O. Palsson, Adil Mardinoglu, Lena Hansson, Mathias Uhlén, and Jens Nielsen. An atlas of human metabolism. Science Signaling, 13(624), March 2020. ISSN 1937-9145. doi: 10.1126/scisignal.aaz1482.

20. Jiahao Luo, Hao Wang, Devlin Moyer, Zhetao Guo, Jonathan L. Robinson, Johan Gustafsson, Mihail Anton, Yu Chen, Eduard J. Kerkhoven, Jens Nielsen, and Feiran Li. Reconstruction of human metabolic models with large language models. Proceedings of the National Academy of Sciences, 123(15), April 2026. ISSN 1091-6490. doi: 10.1073/pnas.2516511123.

21. Stephen Chapman, Theo Brunet, Arnaud Mourier, and Bianca H Habermann. Mitomammal: a genome scale model of mammalian mitochondria predicts cardiac and bat metabolism. Bioinformatics Advances, 5(1), November 2024. ISSN 2635-0041. doi: 10.1093/bioadv/vbae172.

22. Sarah C Dyer, Olanrewaju Austine-Orimoloye, Andrey G Azov, Matthieu Barba, If Barnes, Vianey Paola Barrera-Enriquez, Arne Becker, Ruth Bennett, Martin Beracochea, Andrew Berry, Jyothish Bhai, Simarpreet Kaur Bhurji, Sanjay Boddu, Paulo R Branco Lins, Lucy Brooks, Shashank Budhanuru Ramaraju, Lahcen I Campbell, Manuel Carbajo Martinez, Mehrnaz Charkhchi, Lucas A Cortes, Claire Davidson, Sukanya Denni, Kamalkumar Dodiya, Sarah Donaldson, Bilal El Houdaigui, Tamara El Naboulsi, Oluwadamilare Falola, Reham Fatima, Thiago Genez, Jose Gonzalez Martinez, Tatiana Gurbich, Matthew Hardy, Zoe Hollis, Toby Hunt, Mike Kay, Vinay Kaykala, Diana Lemos, Disha Lodha, Nourhen Mathlouthi, Gabriela Alejandra Merino, Ryan Merritt, Louisse Paola Mirabueno, Aleena Mushtaq, Syed Nakib Hossain, José G Pérez-Silva, Malcolm Perry, Ivana Piližota, Daniel Poppleton, Irina Prosovetskaia, Shriya Raj, Ahamed Imran Abdul Salam, Shradha Saraf, Nuno Saraiva-Agostinho, Swati Sinha, Botond Sipos, Vasily Sitnik, Emily Steed, Marie-Marthe Suner, Likhitha Surapaneni, Kyösti Sutinen, Francesca Floriana Tricomi, Ian Tsang, David Urbina-Gómez, Andres Veidenberg, Thomas A Walsh, Natalie L Willhoft, Jamie Allen, Jorge Alvarez-Jarreta, Marc Chakiachvili, Jitender Cheema, Jorge Batista da Rocha, Nishadi H De Silva, Stefano Giorgetti, Leanne Haggerty, Garth R Ilsley, Jon Keatley, Jane E Loveland, Benjamin Moore, Jonathan M Mudge, Guy Naamati, John Tate, Stephen J Trevanion, Andrea Winterbottom, Bethany Flint, Adam Frankish, Sarah E Hunt, Robert D Finn, Mallory A Freeberg, Peter W Harrison, Fergal J Martin, and Andrew D Yates. Ensembl 2025. Nucleic Acids Research, 53(D1): D948–D957, December 2024. ISSN 1362-4962. doi: 10.1093/nar/gkae1071.

23. Elspeth A. Bruford, Bryony Braschi, Paul Denny, Tamsin E. M. Jones, Ruth L. Seal, and Susan Tweedie. Guidelines for human gene nomenclature. Nature Genetics, 52(8):754–758, August 2020. ISSN 1546-1718. doi: 10.1038/s41588-020-0669-3.

24. Adnan Malik, Muhammad Arsalan, Carlos Moreno, Juan Mosquera, Eloy Félix, Tevfik Kizilören, Venkatesh Muthukrishnan, Barbara Zdrazil, Andrew R Leach, and Noel M O’Boyle. Chebi: re-engineered for a sustainable future. Nucleic Acids Research, 54(D1):D1768–D1778, November 2025. ISSN 1362-4962. doi: 10.1093/nar/gkaf1271.

25. Sébastien Moretti, Anne Niknejad, Marco Pagni, and Florence Mehl. Metanetx: a bridge between metabolic resources for enhanced curation and multi-omics data harmonization. Nucleic Acids Research, 54(D1):D617–D622, November 2025. ISSN 1362-4962. doi: 10.1093/nar/gkaf1286.

26. Alex Thomas, Sorena Rahmanian, Aarash Bordbar, Bernhard Ø. Palsson, and Neema Jamshidi. Network reconstruction of platelet metabolism identifies metabolic signature for aspirin resistance. Scientific Reports, 4(1), January 2014. ISSN 2045-2322. doi: 10.1038/srep03925.

27. Julia Hauenstein, Lisa Jeske, Antje Jäde, Mathias Krull, Katrin Dümmer, Julia Koblitz, Anja Tietz, Dieter Jahn, Lorenz Christian Reimer, and Boyke Bunk. Brenda in 2026: a global core biodata resource for functional enzyme and metabolic data within the dsmz digital diversity. Nucleic Acids Research, 54(D1):D527–D534, November 2025. ISSN 1362-4962. doi: 10.1093/nar/gkaf1113.

28. Ulrike Wittig, Maja Rey, Andreas Weidemann, Renate Kania, and Wolfgang Müller. Sabio-rk: an updated resource for manually curated biochemical reaction kinetics. Nucleic Acids Research, 46 (D1):D656–D660, October 2017. ISSN 1362-4962. doi: 10.1093/nar/gkx1065.

29. Jingnan Huang, Frauke Swieringa, Fiorella A. Solari, Isabella Provenzale, Luigi Grassi, Ilaria De Simone, Constance C. F. M. J. Baaten, Rachel Cavill, Albert Sickmann, Mattia Frontini, and Johan W. M. Heemskerk. Assessment of a complete and classified platelet proteome from genome-wide transcripts of human platelets and megakaryocytes covering platelet functions. Scientific Reports, 11(1), June 2021. ISSN 2045-2322. doi: 10.1038/s41598-021-91661-x.

30. Ranjana Hawaldar and Sadhna Sodani. Study of platelet indices in hyperlipidemia. IP Journal of Diagnostic Pathology and Oncology, 3(4):299–303, December 2020. ISSN 2581-3706. doi: 10.18231/2581-3706.2018.0061.

31. Axel Theorell, Johann F Jadebeck, Katharina Nöh, and Jörg Stelling. Polyround: polytope rounding for random sampling in metabolic networks. Bioinformatics, 38(2):566–567, July 2021. ISSN 1367-4811. doi: 10.1093/bioinformatics/btab552.

32. Richard D Paul, Johann F Jadebeck, Anton Stratmann, Wolfgang Wiechert, and Katharina Nöh. hopsy — a methods marketplace for convex polytope sampling in python. Bioinformatics, 40(7), July 2024. ISSN 1367-4811. doi: 10.1093/bioinformatics/btae430.

33. Hulda S Haraldsdóttir, Ben Cousins, Ines Thiele, Ronan M.T Fleming, and Santosh Vempala. Chrr: coordinate hit-and-run with rounding for uniform sampling of constraint-based models. Bioinformatics, 33(11):1741–1743, January 2017. ISSN 1367-4811. doi: 10.1093/bioinformatics/btx052. doi: 10.48550/ARXIV.1903.08008.

34. Aki Vehtari, Andrew Gelman, Daniel Simpson, Bob Carpenter, and Paul-Christian Bürkner. Rank-normalization, folding, and localization: An improved R for assessing convergence of mcmc. 2019.

35. GitHub. Github. https://github.com/, March 2026. Accessed: 2026-03-06.

36. GitLab. Gitlab. https://about.gitlab.com/, March 2026. Accessed: p2026-03-04.

37. Mojtaba Shahin, Muhammad Ali Babar, and Liming Zhu. Continuous integration, delivery and deployment: A systematic review on approaches, tools, challenges and practices. IEEE Access, 5: 3909–3943, 2017. ISSN 2169-3536. doi: 10.1109/access.2017.2685629.

38. GitLab. Gitlab ci/cd. https://docs.gitlab.com/topics/build_your_application/, March 2026. Accessed: 2026-03-04.

39. GitLab. Gitlab runner. https://docs.gitlab.com/runner/, March 2026. Accessed: 2026-03-04.

40. Christian Lieven, Moritz E. Beber, Brett G. Olivier, Frank T. Bergmann, Meric Ataman, Parizad Babaei, Jennifer A. Bartell, Lars M. Blank, Siddharth Chauhan, Kevin Correia, Christian Diener, Andreas Dräger, Birgitta E. Ebert, Janaka N. Edirisinghe, José P. Faria, Adam M. Feist, Georgios Fengos, Ronan M. T. Fleming, Beatriz García-Jiménez, Vassily Hatzimanikatis, Wout van Helvoirt, Christopher S. Henry, Henning Hermjakob, Markus J. Herrgård, Ali Kaafarani, Hyun Uk Kim, Zachary King, Steffen Klamt, Edda Klipp, Jasper J. Koehorst, Matthias König, Meiyappan Lakshmanan, Dong-Yup Lee, Sang Yup Lee, Sunjae Lee, Nathan E. Lewis, Filipe Liu, Hongwu Ma, Daniel Machado, Radhakrishnan Mahadevan, Paulo Maia, Adil Mardinoglu, Gregory L. Medlock, Jonathan M. Monk, Jens Nielsen, Lars Keld Nielsen, Juan Nogales, Intawat Nookaew, Bernhard O. Palsson, Jason A. Papin, Kiran R. Patil, Mark Poolman, Nathan D. Price, Osbaldo Resendis-Antonio, Anne Richelle, Isabel Rocha, Benjamín J. Sánchez, Peter J. Schaap, Rahuman S. Malik Sheriff, Saeed Shoaie, Nikolaus Sonnenschein, Bas Teusink, Paulo Vilaça, Jon Olav Vik, Judith A. H. Wodke, Joana C. Xavier, Qianqian Yuan, Maksim Zakhartsev, and Cheng Zhang. Memote for standardized genome-scale metabolic model testing. Nature Biotechnology, 38(3):272–276, March 2020. ISSN 1546-1696. doi: 10.1038/s41587-020-0446-y.

41. Shona A. Mookerjee, Akos A. Gerencser, David G. Nicholls, and Martin D. Brand. Quantifying intracellular rates of glycolytic and oxidative atp production and consumption using extracellular flux measurements. Journal of Biological Chemistry, 292(17):7189–7207, April 2017. ISSN 0021-9258. doi: 10.1074/jbc.m116.774471.

42. Iván Domenzain, Benjamín Sánchez, Mihail Anton, Eduard J. Kerkhoven, Aarón Millán-Oropeza, Céline Henry, Verena Siewers, John P. Morrissey, Nikolaus Sonnenschein, and Jens Nielsen. Reconstruction of a catalogue of genome-scale metabolic models with enzymatic constraints using gecko 2.0. Nature Communications, 13(1), June 2022. ISSN 2041-1723. doi: 10.1038/s41467-022-31421-1.

43. Jun Fan, Changliang Shan, Hee-Bum Kang, Shannon Elf, Jianxin Xie, Meghan Tucker, Ting-Lei Gu, Mike Aguiar, Scott Lonning, Huaibin Chen, Moosa Mohammadi, Laura-Mae P. Britton, Benjamin A. Garcia, Maša Alečković, Yibin Kang, Stefan Kaluz, Narra Devi, Erwin G. Van Meir, Taro Hitosugi, Jae Ho Seo, Sagar Lonial, Manila Gaddh, Martha Arellano, Hanna J. Khoury, Fadlo R. Khuri, Titus J. Boggon, Sumin Kang, and Jing Chen. Tyr phosphorylation of pdp1 toggles recruitment between acat1 and sirt3 to regulate the pyruvate dehydrogenase complex. Molecular Cell, 53(4):534–548, February 2014. ISSN 1097-2765. doi: 10.1016/j.molcel.2013.12.026.

44. Brandon R Desousa, Kristen KO Kim, Anthony E Jones, Andréa B Ball, Wei Y Hsieh, Pamela Swain, Danielle H Morrow, Alexandra J Brownstein, David A Ferrick, Orian S Shirihai, Andrew Neilson, David A Nathanson, George W Rogers, Brian P Dranka, Anne N Murphy, Charles Affourtit, Steven J Bensinger, Linsey Stiles, Natalia Romero, and Ajit S Divakaruni. Calculation of atp production rates using the seahorse xf analyzer. The EMBO Reports, 24(10), August 2023. ISSN 1469-3178. doi: 10.15252/embr.202256380.

45. Irina Portier and Matthew T. Rondina. Platelets, plasma, and proteostasis: a translation tightrope. Blood Advances, 8(6):1567–1569, March 2024. ISSN 2473-9537. doi: 10.1182/bloodadvances.2023012401.

46. Bingsen Zhang and Frank C. Schroeder. Mechanisms of metabolism-coupled protein modifications. Nature Chemical Biology, 21(6):819–830, January 2025. ISSN 1552-4469. doi: 10.1038/s41589-024-01805-z.

47. Yu Chen, Johan Gustafsson, Albert Tafur Rangel, Mihail Anton, Iván Domenzain, Cheewin Kittikunapong, Feiran Li, L. Yuan Jens Nielsen, and Eduard J. Kerkhoven. Reconstruction, simulation and analysis of enzyme-constrained metabolic models using gecko toolbox 3.0. Nature Protocols, 19(3):629–667, January 2024. ISSN 1750-2799. doi: 10.1038/s41596-023-00931-7.

48. Jorge Carrasco Muriel, Christopher Long, and Nikolaus Sonnenschein. Simultaneous application of enzyme and thermodynamic constraints to metabolic models using an updated python implementation of gecko. Microbiology Spectrum, 11(6), December 2023. ISSN 2165-0497. doi: 10.1128/spectrum.01705-23.

49. Sunghwan Kim, Jie Chen, Tiejun Cheng, Asta Gindulyte, Jia He, Siqian He, Qingliang Li, Benjamin A Shoemaker, Paul A Thiessen, Bo Yu, Leonid Zaslavsky, Jian Zhang, and Evan E Bolton. Pubchem 2025 update. Nucleic Acids Research, 53(D1):D1516–D1525, November 2024. ISSN 1362-4962. doi: 10.1093/nar/gkae1059.

