## supplementary information for "Curating MitoCore: A Standardized Small-Scale Human Metabolic Model as Platform for Proteomics Integration and Disease Modeling"

#### 1 A new Algorithm for autoPACMEN's Calibration Candidate selection and extension of the Calibrator

AutoPACMEN is a library for Python and Matlab for incorporating quantitative proteomics data into metabolic models by the sMOMENT method (Bekiaris and Klamt, 2020). Essentially, sMOMENT introduces an upper limit to reactions determined from the product of available enzyme concentration and the reaction's k_cat value. This enzyme resource limit is introduced to reactions through pseudo-metabolites representing the pool of available protein resources. autoPACMEN introduces two types of pseudo-metabolites: A global protein pool representing all protein resources of a cell and pools for individual proteins measured by proteomics. It is possible to constrain the model exclusively through the global protein pool if no individual protein concentrations are available.

AutoPACMEN consists of a "model generator" fetching k_cat values from BRENDA (Hauenstein, et al., 2025) and Sabio-RK (Wittig, et al., 2017) introducing the enzyme constraints to an input model and the "model calibrator" fitting the FBA objective function values to measured fluxes by altering k_cat values. The fitting step is typically necessary because the objective value cannot be reached by uncalibrated sMOMENT models due to restrictive k_cat values. This usually only applies to a fraction of all model reactions, requiring only k_cats of these reactions to be calibrated, which could be determined by a dedicated Python function of the "generator". Originally, this function did only work for reactions coupled to the global protein pool, but not if pools for individual proteins (enzymes) were introduced.

We implemented an algorithm for determining candidate reactions for calibration based on sensitivity $C_{R_i}^{Z}$ of the objective function $Z$ to the turnover number of each individual reaction $k_{cat,R_i}$, similar to the concept of the flux control coefficient (Burns, et al., 1985):

$$C_{R_i}^{Z} = \frac{dZ}{dk_{cat,R_i}} \frac{k_{cat,R_i}}{Z} \quad (1)$$

Reactions with sensitivities above a defined threshold (default 0.001) are selected as candidates. k_cats of these candidates are increased by ten-fold to relax the enzyme constraints. This process is repeated several times until the target objective value can be reached or if the maximal number of iterations (default 5 iterations) is met. The candidates from every iteration are subjected to the "calibrator" which was also adapted to consider individual protein pools during the calibration. The output of our calibration script are factors that are multiplied with the original k_cats to yield the calibrated parameters.

The calibrator module of autoPACMEN is intended to fit kcat values of selected reactions to measured fluxes. Our script (modelcalibration_05.m) reads candidate reactions for calibration (".\autopacmen_output\kcat_candidates.json") and scenarios (".\model_data_input\platelet_scenario.json") from .json files prepared beforehand by the jupyter notebooks.

Calibration is performed for each model individually first calibrating kcat values and subsequently calibrating the stoichiometry of the protein pool delivery reaction. This order is chosen because kcats coupled to individual protein pools are more restrictive than the total protein pool and the objective is lower than the target objective.

AutoPACMEN was originally only calibrating the stoichiometric coefficients M/kcat of the global pool metabolite (M_prot_pool) in reactions and did not consider pools for individual proteins and their stoichiometric coefficients (1/kcat). Instead of changing the stoichiometric coefficients directly during the calibration, we introduced a factor for each kcat value (start_kcat_factors) that scales the stoichiometric coefficients of pseudo-metabolites. Effectively, this factor is calibrated. The advantage here is that all stoichiometric coefficients of pseudo-metabolites are scaled at the same time and there is no need to keep track of them separately (they are all based on the kcat). The same kcats are assigned to parallel reactions catalyzed by isoenzyme splits of the original reaction.

To minimize arbitrary combinations of kcats that support the target value, a second optimization round was introduced, minimizing the deviation of kcats to their original values. This step is analogously to pFBA where the total sum of fluxes is minimized after the objective has been calculated to remove unnecessarily high fluxes (Lewis, et al., 2010).

After kcats have been calibrated, the lowest protein pool still supporting the target objective is determined by line search. The resulting calibrated protein pool represents the minimal global protein resources necessary to meet the target reaction flux.

The source code of all changed autoPACMEN functions is included in the source code repository (https://doi.org/10.5281/zenodo.20813825).

#### 2 Evaluation of Prediction Errors

Model predictions from flux sampling were evaluated by comparing the error between prediction and measured flux data from (Sake, et al., 2022) and Seahorse analysis (JedliČKa, et al., 2021). Because flux sampling distributions are not necessarily normally distributed, we reported the median and limits of the interquartile range (IQR, 25% and 75% quantiles, see tables 2 and 3 in the main manuscript). Evaluations were done for exchange fluxes of glucose, acetate and lactate, as well as for the two main origins of ATP, glycolysis and oxidative phosphorylation (OXPHOS).

To evaluate how well the most likely measurement aligns with the most likely prediction, we determined the root mean squared relative error (RMSRE) between medians of measurements ($r_i$) and predictions ($\hat{r}_i$):

$$RMSRE = \sqrt{\frac{1}{n}\sum_{i=1}^{n}\frac{(r_i - \hat{r}_i)^2}{r_i}} \tag{2}$$

Instead of using the residual as in the root mean squared error, equation 2 uses the relative error to account for different flux magnitudes between reactions.

Furthermore, we evaluated whether predicted flux distributions fall into a biologically feasible range. To this end, standard deviations ($s$) of the measured reactions ($i$) were transformed into 95% confidence intervals ($CI$) considering the critical value from the t-statistic ($t_{crit}$, determined for two-tailed confidence level of 0.05 and N-1 degrees of freedom) and the number of biological samples $N$:

$$CI_i = r_i \pm t_{crit}\frac{s_i}{\sqrt{N}} \tag{3}$$

Based on these confidence intervals, we determined the mean coverage probability (MCP) as number of flux samples falling into the 95% confidence range for each reaction ($n_{i,s,CI}$) divided by the total number of flux samples ($n_{i,s}$), averaged over all $n$ measured reactions:

$$MCP = \frac{1}{n}\sum_{i=1}^{n}\frac{n_{i,s,CI}}{n_{i,s}} \tag{4}$$

If all flux samples fall into the 95% confidence intervals, the MCP assumes 1.0 and it becomes 0.0, if no samples are in this range. Unfortunately, no information on the number of biological replicates for the calculation of exchange fluxes was given in(Sake, et al., 2022), therefore it was not possible to determine the MCP for these measurements.

#### 3 Error Propagation of Seahorse Standard deviation

The fraction of ATP produced by glycolysis (f_glyco) was calculated from the given Seahorse net fluxes for glycolysis (r_glyco) and oxidative phosphorylation (r_oxphos):

$$f_{glyco} = \frac{r_{glyco}}{r_{glyco} + r_{oxphos}} \tag{5}$$

To determine the standard deviation of f_glyco (s_f,glyco), error propagation provides the following formula utilizing the given standard deviation of the glycolysis net flux (s_glyco) and oxphos net flux (s_oxphos):

$$s_{f,glyco}^2 = \left(\frac{\partial f_{glyco}}{\partial r_{glyco}}\right)^2 s_{glyco}^2 + \left(\frac{\partial f_{glyco}}{\partial r_{oxphos}}\right)^2 s_{oxphos}^2 \tag{6}$$

Equation 6 can be resolved to:

$$s_{f,glyco} = \frac{\sqrt{r_{oxphos}^2 s_{glyco}^2 + r_{glyco}^2 s_{oxphos}^2}}{\left(r_{glyco} + r_{oxphos}\right)^2} \tag{7}$$

### 4 Full Comparison of MitoCore and Human-GEM

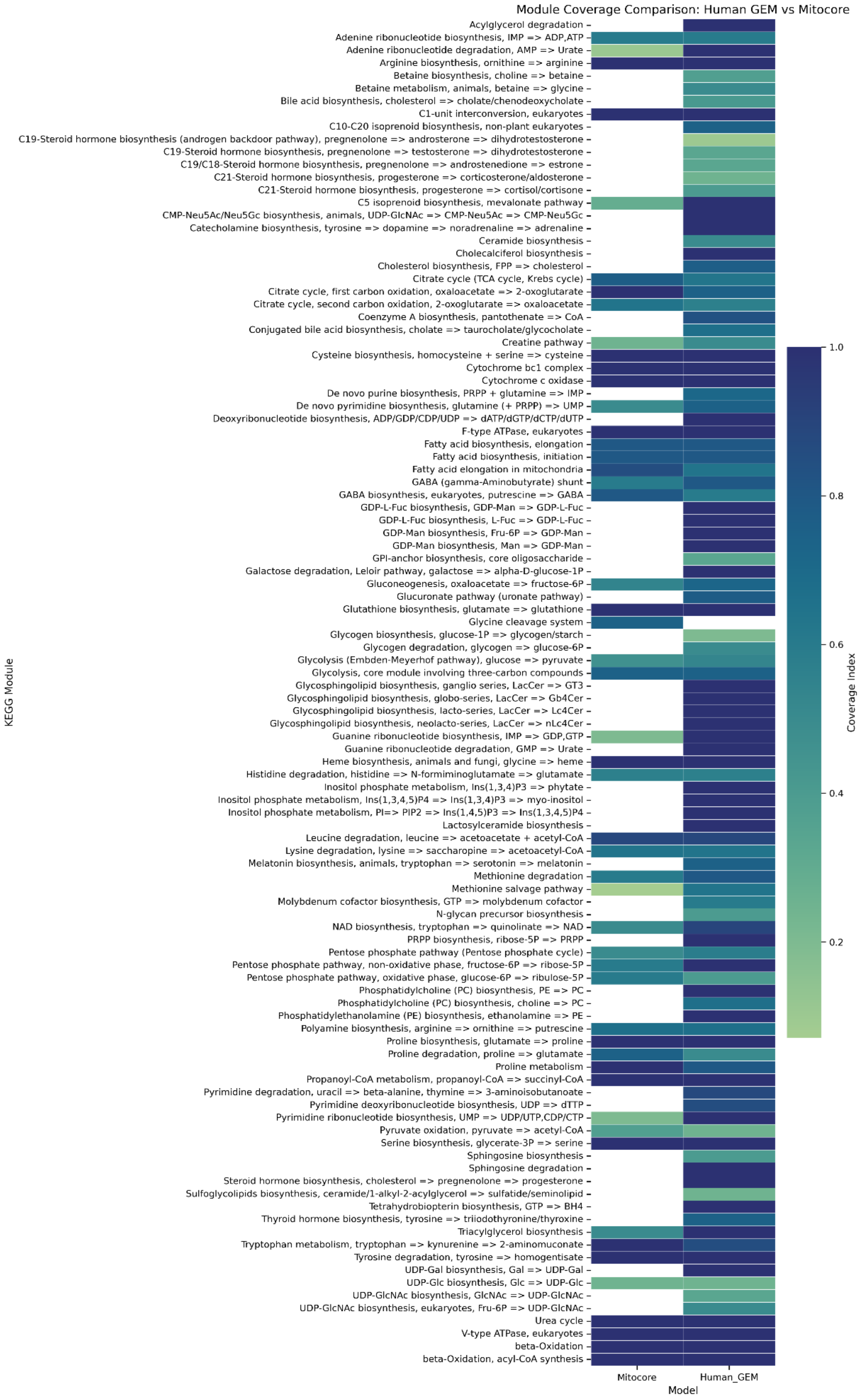


*Figure S1 Coverage of KEGG modules by reactions in MitoCore and Human-GEM. Numerical coverage values can be reviewed in the file "model_curation\Pathway_Coverage_Files\coverage_index_modules.xlsx" included in the source code repository (https://doi.org/10.5281/zenodo.20813825)*

#### 5 Flux Distributions for all exchange reactions

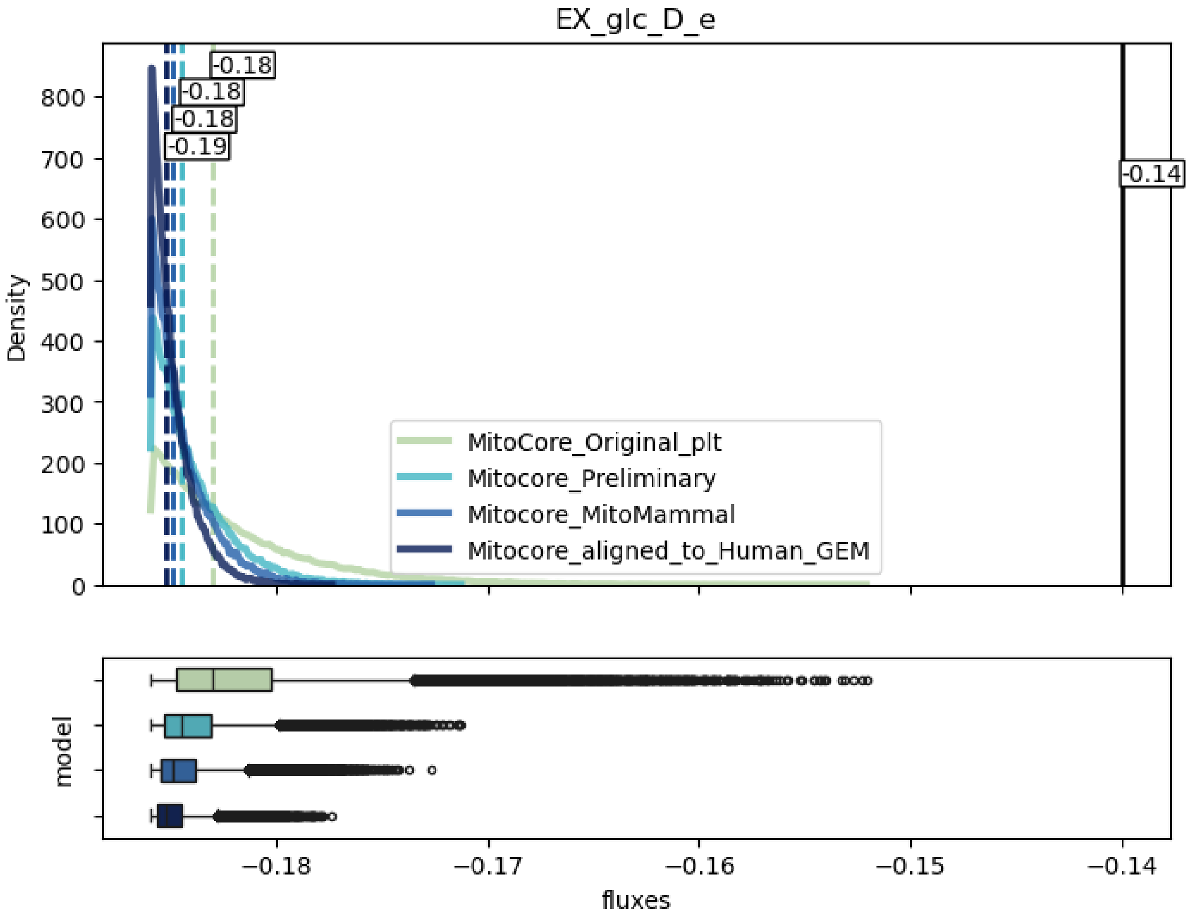


*Figure S2 Predicted flux distributions for glucose uptake by flux sampling. Shown distributions support at least 95% of the FBA optimum of total ATP production. The black dashed line represents the measured glucose uptake from 13C experiments (Sake, et al., 2022). Colored dashed lines represent the median fluxes for flux distributions. The boxplots below the density plots represent the same flux sampling distributions for each model as the density plot.*

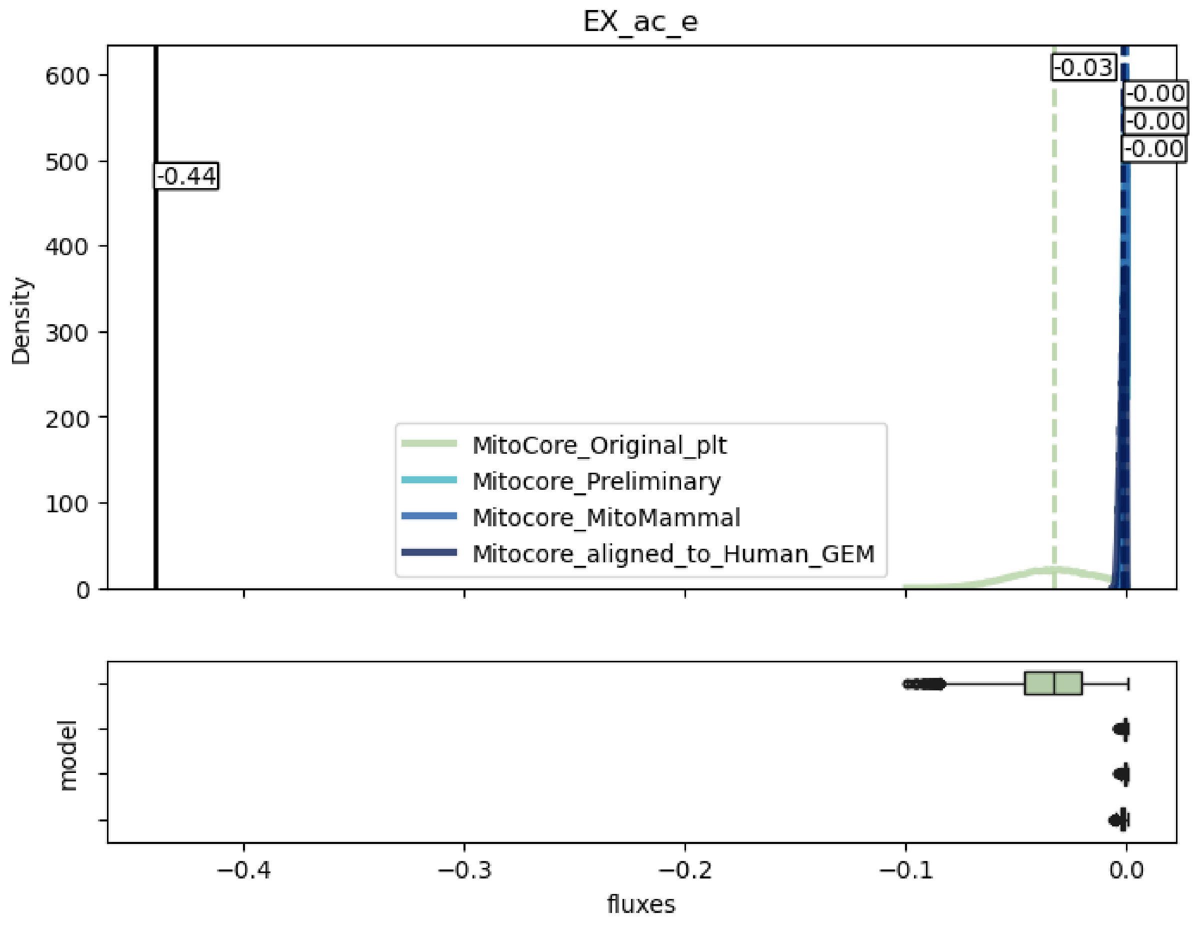


*Figure S3 Predicted flux distributions for acetate uptake by flux sampling. Shown distributions support at least 95% of the FBA optimum of total ATP production. The black dashed line represents the measured glucose uptake from 13C experiments (Sake, et al., 2022). Colored dashed lines represent the median fluxes for flux distributions. The boxplots below the density plots represent the same flux sampling distributions for each model as the density plot.*

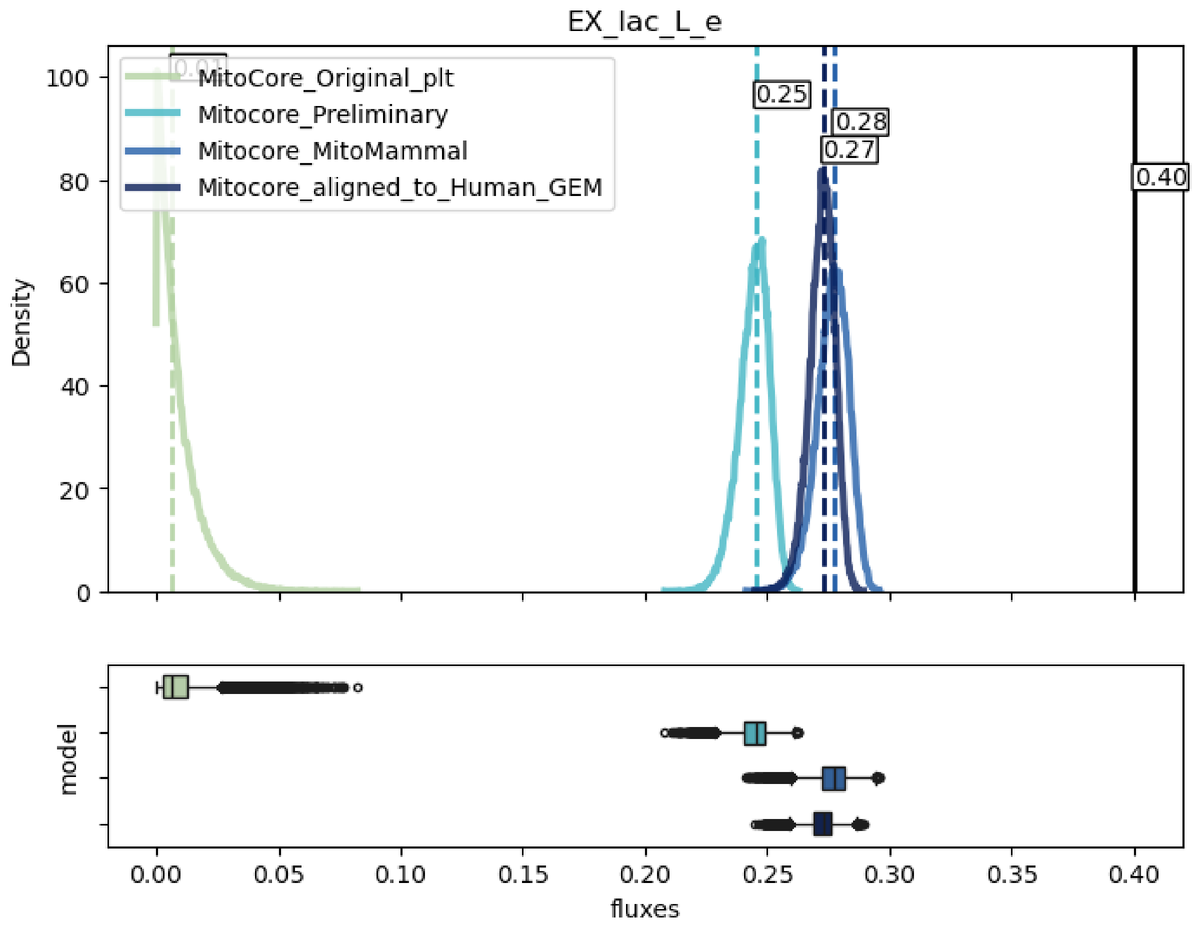


*Figure S4 Predicted flux distributions for lactate secretion by flux sampling. Shown distributions support at least 95% of the FBA optimum of total ATP production. The black dashed line represents the measured glucose uptake from 13C experiments (Sake, et al., 2022). Colored dashed lines represent the median fluxes for flux distributions. The boxplots below the density plots represent the same flux sampling distributions for each model as the density plot.*

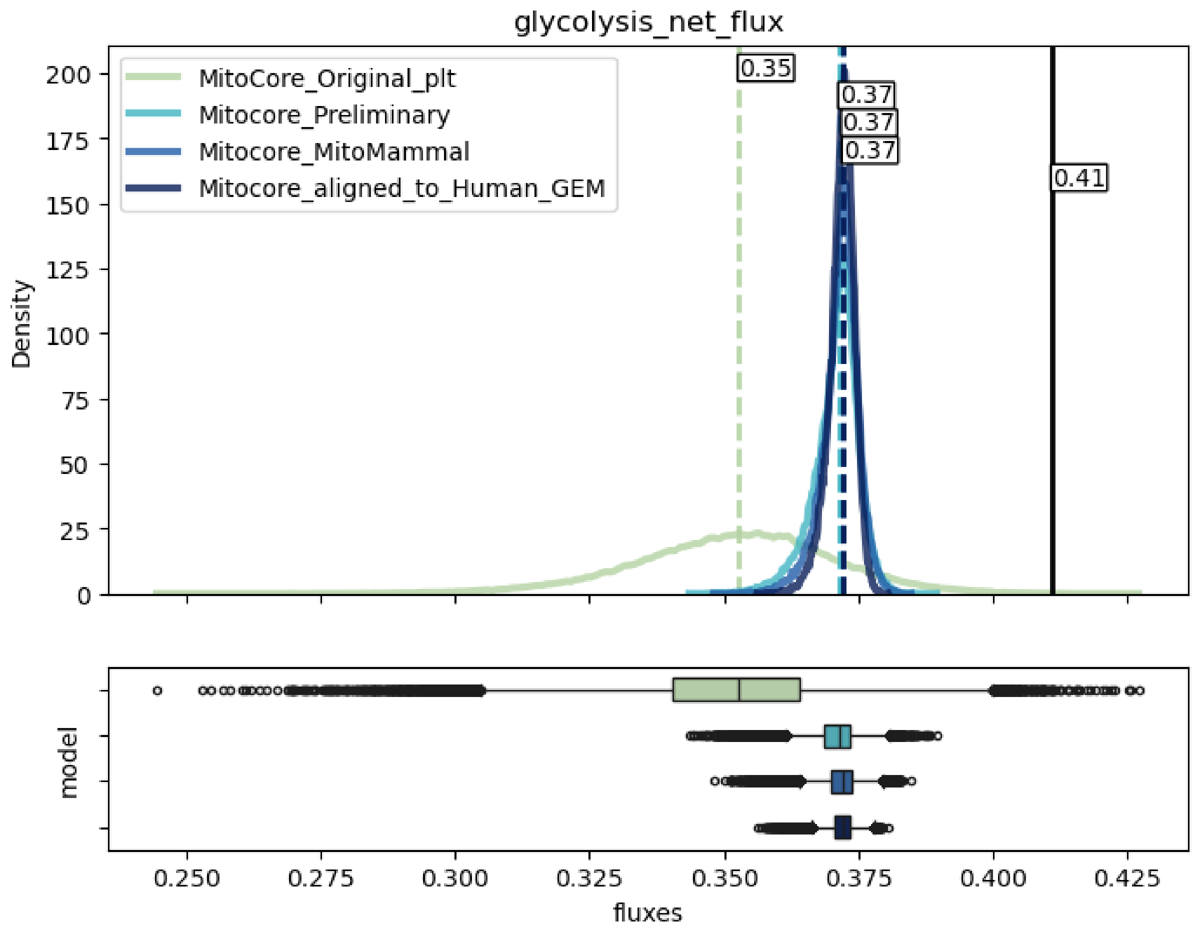


*Figure S5 Predicted flux distributions for glycolysis net ATP production by flux sampling. Glycolysis net flux was calculated from fluxes of Pyruvate Kinase (PYK, r0122), Phosphoglycerate kinase 1 (PGK), Hexokinase (HEX1), and Phosphofructokinase (PFK). Shown distributions support at least 95% of the FBA optimum of total ATP production. The black dashed line represents the measured glucose uptake from 13C experiments (Sake, et al., 2022). Colored dashed lines represent the median fluxes for flux distributions. The boxplots below the density plots represent the same flux sampling distributions for each model as the density plot.*

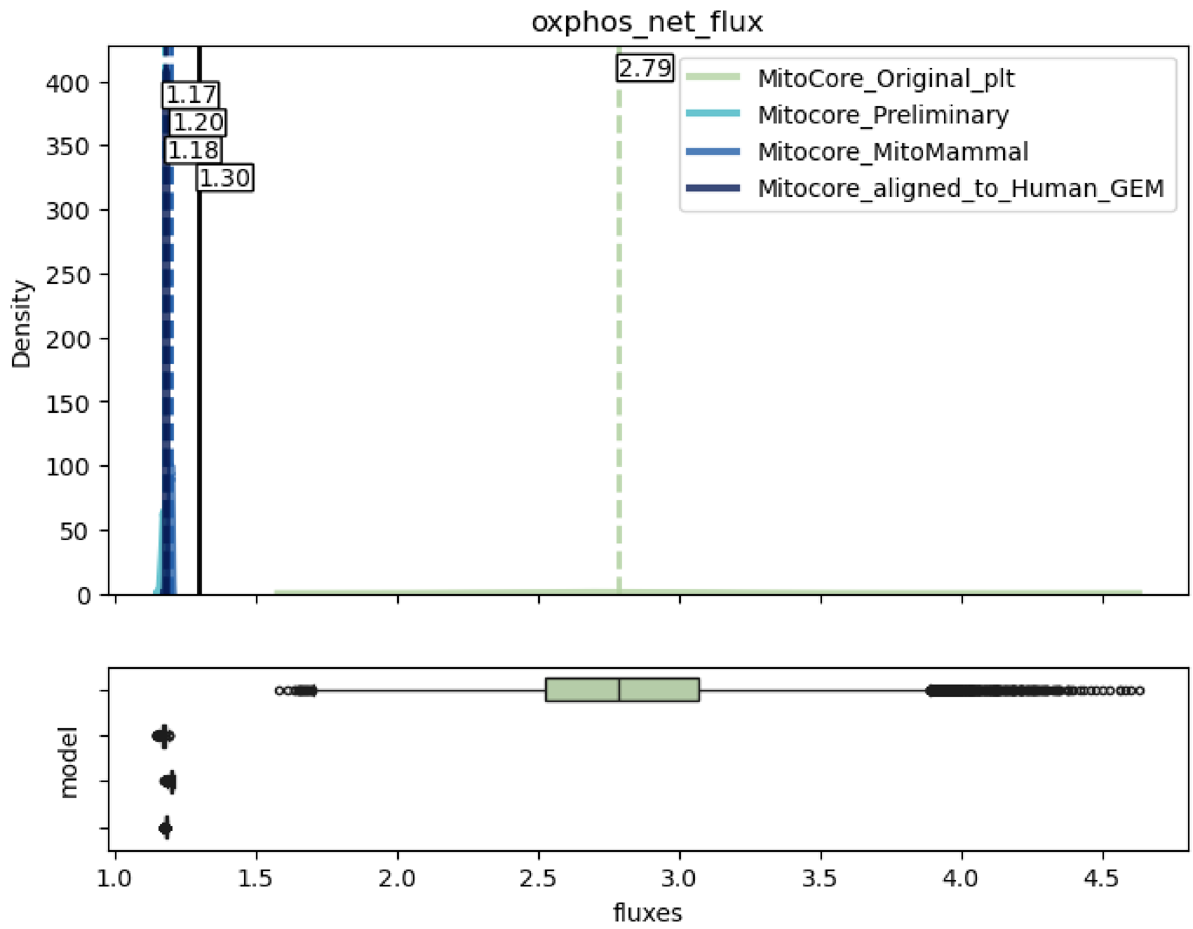


*Figure S6 Predicted flux distributions for OXPHOS net ATP production by flux sampling. OXPHOS net ATP flux was determined by the flux of ATP synthase (CV_MitoCore). Shown distributions support at least 95% of the FBA optimum of total ATP production. The black dashed line represents the measured glucose uptake from 13C experiments (Sake, et al., 2022). Colored dashed lines represent the median fluxes for flux distributions. The boxplots below the density plots represent the same flux sampling distributions for each model as the density plot.*

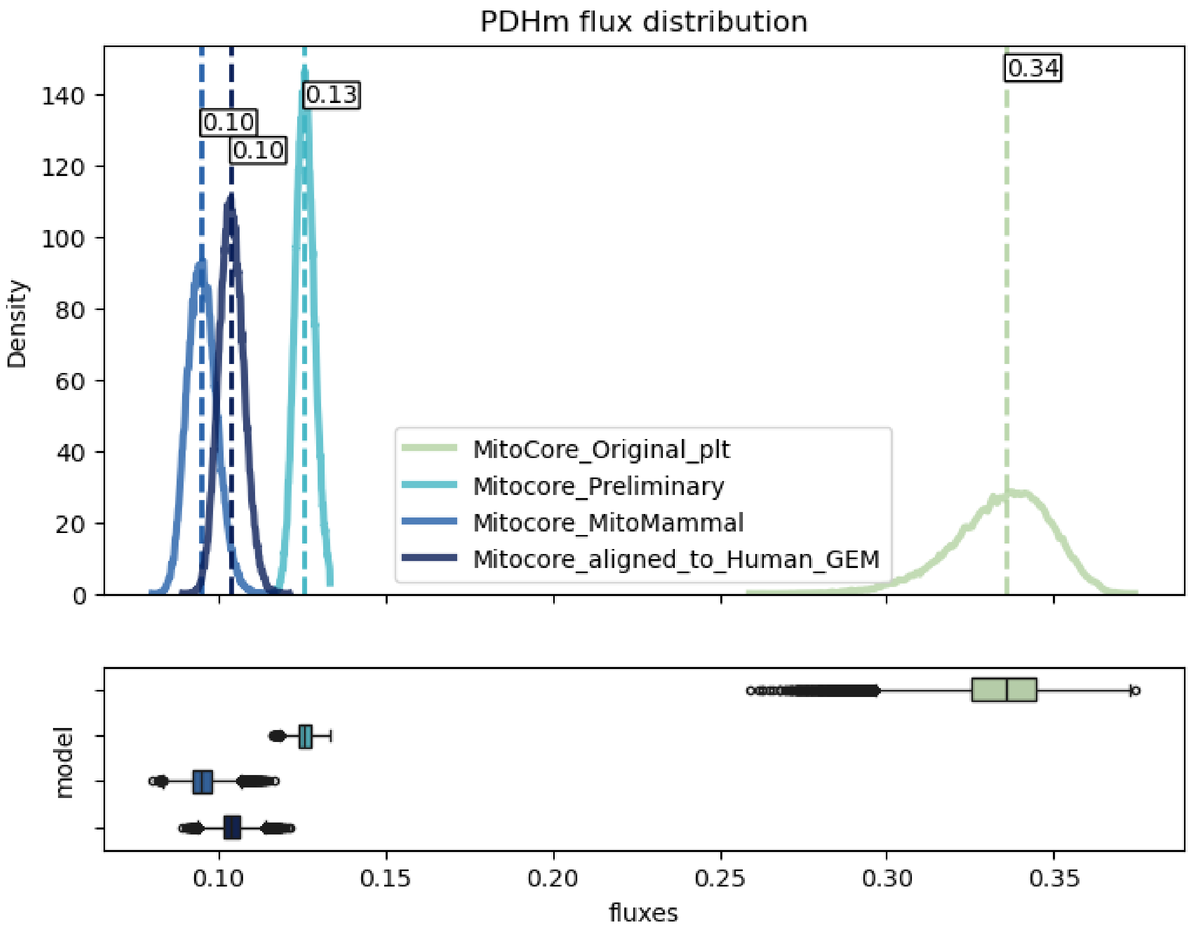


*Figure S7 Predicted flux distributions for PDHm (PDHm_GPRSPLIT_1 in MitoCore Preliminary and MitoMammal, armr_PDHm in MitoCore Human-GEM) by flux sampling. Shown distributions support at least 95% of the FBA optimum of total ATP production. Colored dashed lines represent the median fluxes for flux distributions. The boxplots below the density plots represent the same flux sampling distributions for each model as the density plot.*

### 6 Detailed Curation Pipeline Overview

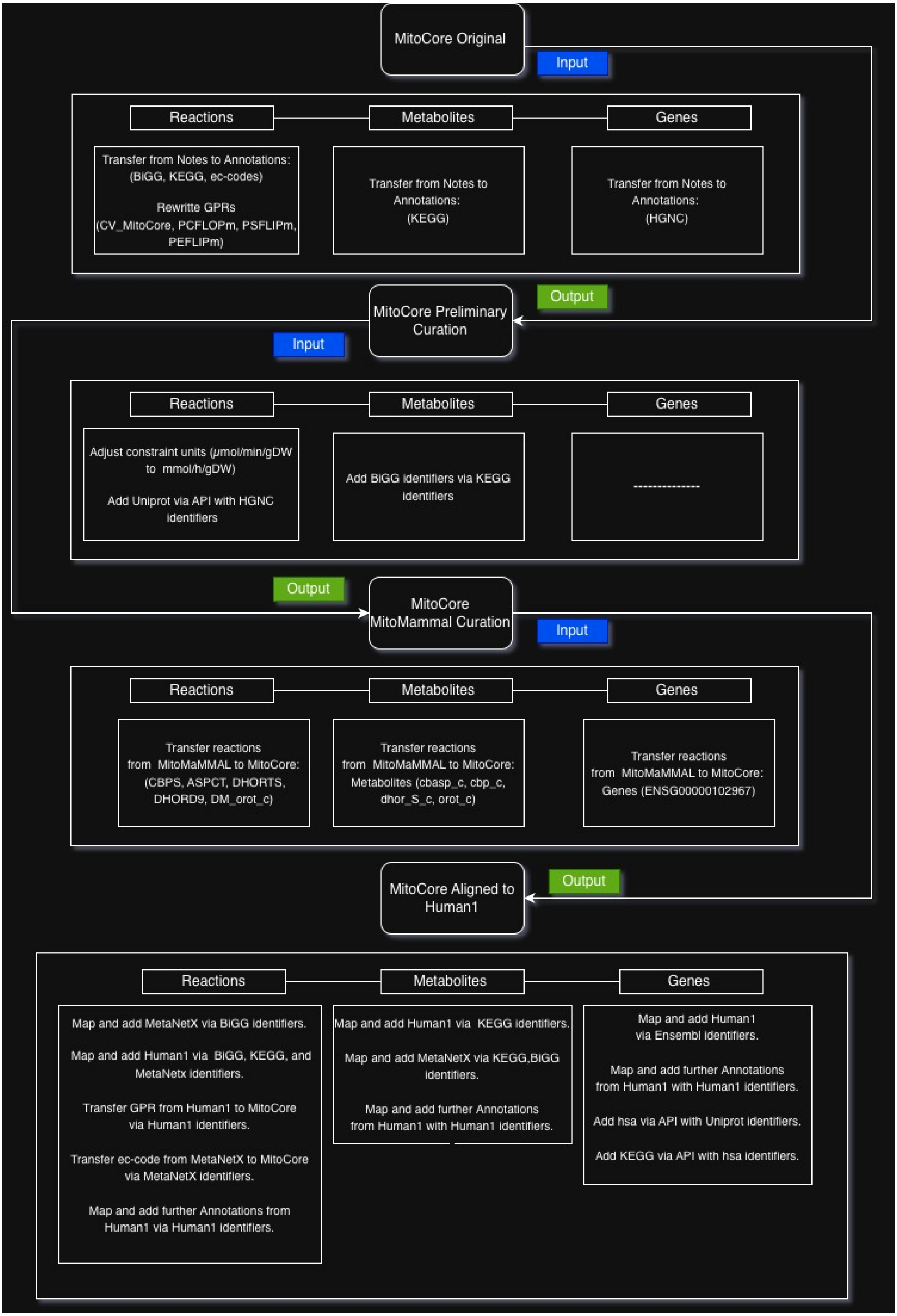


*Figure S7: Schematic overview of all curation steps.*

#### References


Bekiaris, P.S. and Klamt, S. Automatic construction of metabolic models with enzyme constraints. *BMC Bioinformatics* 2020;21(1).

Burns, J.A., *et al*. Control analysis of metabolic systems. *Trends in Biochemical Sciences* 1985;10(1):16-16.

Hauenstein, J., *et al*. BRENDA in 2026: a Global Core Biodata Resource for functional enzyme and metabolic data within the DSMZ Digital Diversity. *Nucleic Acids Research* 2025;54(D1):D527–D534-D527–D534.

JedliČKa, J., R, K.U.N.C. and KuncovÁ, J. Mitochondrial Respiration of Human Platelets in Young Adult and Advanced Age – Seahorse or O2k? *Physiological Research* 2021:S369–S379-S369–S379.

Lewis, N.E., *et al*. Omic data from evolved E. coli are consistent with computed optimal growth from genome-scale models. *Molecular Systems Biology* 2010;6(1).

Sake, C.L., *et al*. Isotopically nonstationary 13C metabolic flux analysis in resting and activated human platelets. *Metabolic Engineering* 2022;69:313–322-313–322.

Wittig, U., *et al*. SABIO-RK: an updated resource for manually curated biochemical reaction kinetics. *Nucleic Acids Research* 2017;46(D1):D656–D660-D656–D660.
